# Capturing rapid learning in an extended successor representation theory of the cognitive map

**DOI:** 10.64898/2025.12.25.696522

**Authors:** Suhee Cho, James L. McClelland

## Abstract

Animals sometimes adapt their behavior after a single exposure to novel information. Here, we integrate abstract theory with recent mechanistic insights to understand this and other examples of rapid adaptation. We ground our investigation in studies of animal learning in spatial contexts, where it is widely held that learning depends on the formation of a cognitive map reflected in learned representations in hippocampal areas CA1 and CA3. We extend the successor representation (SR) theory of the cognitive map, incorporating the weighting of this representation by perceived salience (PS) signals, and using it to represent future availability of environmental features that allow behavior to be guided flexibly by an animal’s goals and needs at later times. To capture rapid adaptation, we rely on behavioral time-scale synaptic plasticity (BTSP), a form of plasticity that captures one-shot formation of new place fields. Through simulations, we show that our hippocampal model network can learn the PS-weighted SR rapidly and set up the ability to replay experiences in the environment after a single exposure through BTSP. Spontaneous replay activity during subsequent offline periods strengthens the PS-weighted SR. Predictions of environmental features also develop rapidly, and behavioral simulations show how these learned representations give rise to need-dependent choice, rapid behavioral adaptation to changes in the availability of reward, and one-trial shock avoidance learning.

## Introduction

Animals’ behavior often adapts quickly to experience. How is this rapid adaptation achieved? This question is important for both neuroscience and artificial intelligence (AI). Many existing models in both computational neuroscience and AI can capture aspects of learned human and animal abilities, but require orders of magnitude more experience to learn these abilities than humans or animals do. Understanding how the brain can learn so efficiently would therefore address a key limitation in our current understanding of learning both in ourselves and our machines.

Here, we ground our exploration of this question within the context of studies of animals behaving in spatial environments (Fig. 1a). In such settings, animals can show very rapid adaptation, often altering their behavior after a single exposure to a salient environmental feature^1^. We further situate our work in the framework of the “cognitive map,” a hypothetical construct that receives locally available information, integrates it with internal state information, and generates signals that guide behavior in spatial environments^2–4^. Recording studies in the hippocampus of behaving animals revealed that the existence of place cells collectively cover all of the locations within a spatial arena^5^, such that they can be thought of as a central component of a cognitive map^6^. Furthermore, experiments show that rodents form such a map rapidly, with many new place fields forming during an initial traversal of a novel environment^7^. Our work aims to provide an integrative framework for understanding how this rapid and efficient representation learning comes about and how it then influences behavior.

**Figure 1.**
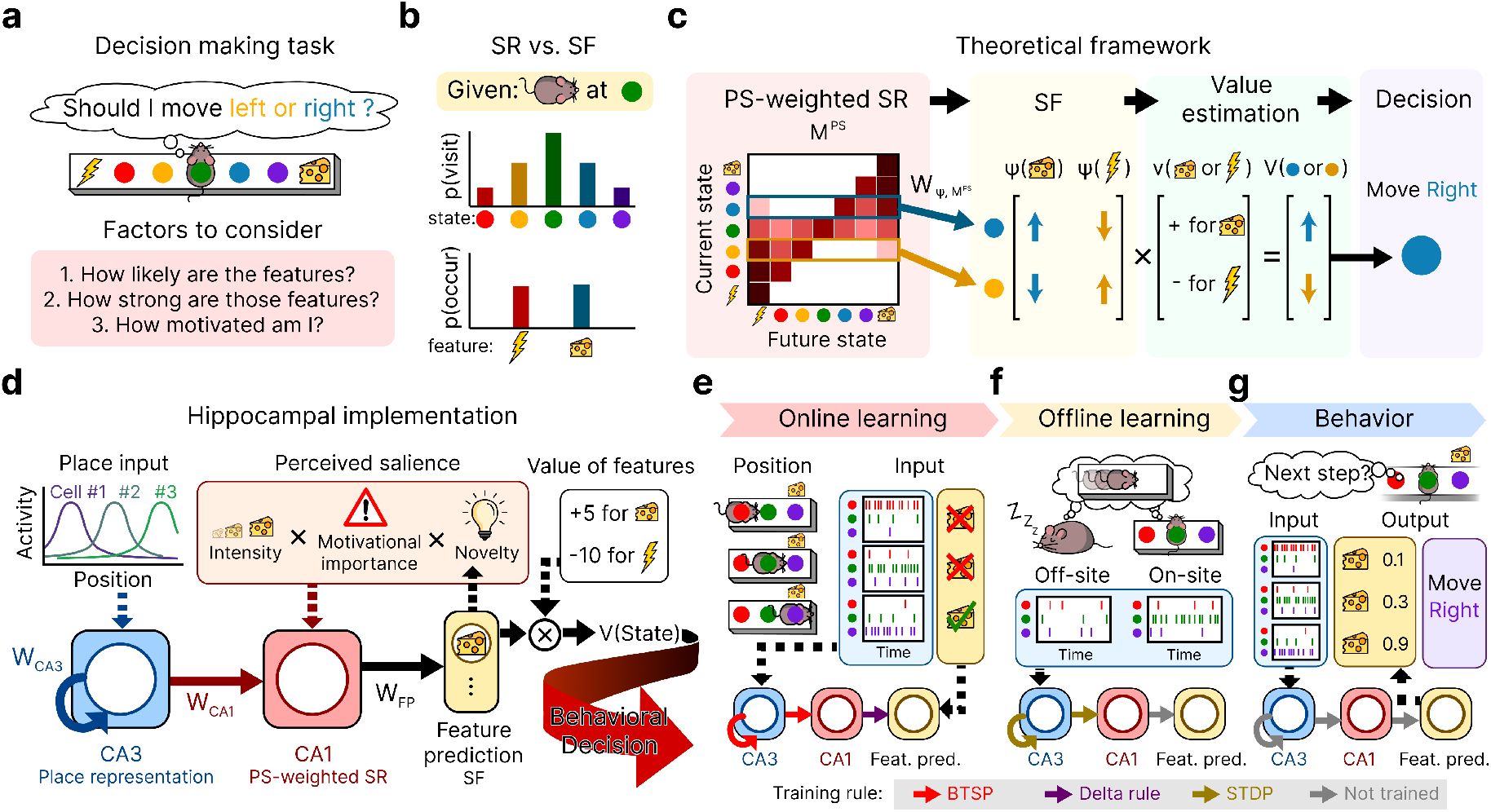
**a**, Rodent decision-making in a maze. **b**, Two ways of representing a state: successor representation (SR) and successor features (SF). **c**, Schematic of the theoretical framework. A PS–weighted SR is relayed to predict successor features, which are then used to compute value and guide decisions. **d**, Hippocampal implementation of the model. CA3 receives place input and encodes place representations. CA1 receives input from CA3 along with P, and encodes the PS-weighted SR. This is passed to the feature prediction layer, which encodes SFs and is ultimately used to determine the subjective value of states. **e–g**, Three distinct phases of our simulations. **e**, *Online learning*: the animal actively explores the environment and encounters features. Place input varies with position, and hippocampal weights are updated using the BTSP rule, while the feature-prediction weights are trained using an error-correcting delta rule. **f**, *Offline learning*: simulating rest or sleep. Place input is held constant, and hippocampal weights are updated using symmetric STDP while the feature-prediction weights are fixed. **g**, *Behavior simulation*: Value-based decision making in the learned track. No weights are updated. The learned connection weights are used to calculate feature predictions, which are assigned value based on current motivational state to guide behavior toward higher-valued outcomes.

We build on the seminal account of the cognitive map provided by successor representation (SR) theory^8^ and its application to hippocampal representation learning^9^. Rather than encoding environments purely in terms of spatial distance, the SR encodes the expected future occupancy of each location based on the animal’s movement history (Fig. 1b, top). Although this approach is supported by experimentally observed properties of hippocampal place cells, including the predictive skewing of place fields^10,11^, the original version of this theory^9^ has two important limitations.

First, as originally described^9^, this account does not explain how factors such as reward^12^, aversive outcomes^1^, or novelty^13^ affect hippocampal representations. These and other factors, collectively termed perceived salience (PS)^14^, can modulate hippocampal representations, including increases in the number of place cells encoding salient locations^12,15–18^ and accelerated formation of those fields^19^. Together, these findings suggest that the hippocampus integrates PS signals with spatial inputs, producing a PS-weighted SR.

Second, the original SR theory does not address how rapid representational adaptation is achieved. This may reflect the framing of the approach as a normative or computational level^20^ account, where the priority has been to capture emergent functional characteristics that the SR enables rather than speed of learning *per se*, and in part from the reliance on learning rules available at the time the work was done. In any case, past simulations showing how the SR may be acquired^9,21^ typically rely on reinforcement-learning-theory based learning rules^22^ or longstanding biologically proposed plasticity rules^23^, both of which can require many (up to thousands) of traversals through an environment to learn.

The discovery of behavioral time-scale synaptic plasticity (BTSP) provides a candidate mechanism that makes it possible to work toward treating rapid adaptation as an important functional characteristic to be addressed. First described nearly a decade ago^12^, BTSP occurs when neurons undergo dendritic plateau potentials that produce large synaptic changes linking neural activity across a time scale of seconds, leading to one-trial formation of new place fields^12,24^. BTSP is coming to be recognized as a potentially viable basis for rapid neural adaptation^25,26^, capturing formation and change of place fields better than alternative spike-time dependent learning rules, which are too weak to capture these changes within realistic parameter ranges^27^. In hippocampal area CA1, BTSP creates place fields that exhibit SR-like anticipatory predictions of future states^12^ and has recently been used to model rapid formation of the classical (non–PS-weighted) SR and reward-directed navigation^28^. Here, we extend this approach to incorporate PS-weighting of predictive representations in CA1, consistent with findings that BTSP in this region is modulated by novelty, stimulus intensity, and reward^19,29–31^.

Our work further highlights how BTSP can initiate a virtuous cycle of adaptation by enabling hippocampal replay after a single exposure to a novel environment or a motivationally salient event. Hippocampal replay was first observed during periods of sleep^32^ and identified as occurring during bursts of activity during which hippocampal place cells spontaneously reactivate in patterns that recapitulate trajectories through an environment^33^. This phenomenon has long been thought to contribute to the strengthening, stabilization, and/or consolidation of learning acquired during physical exploration of the environment^34–37^. More recent studies show evidence of replay during waking behavior, sometimes while animals are at rest after consuming a reward^38^, and sometimes during or prior to exploration^1,39^. We rely on BTSP in hippocampal area CA3 to rapidly create connections among CA3 place cells to enable replay events that occur in subsequent offline periods^27,40,41^. Remarkably, these sequences can represent trajectories that the animal has never physically traversed^42^, suggesting that replay helps complete the internal spatial map^7,43^, and we demonstrate how this can occur through further synaptic plasticity thought to occur during replay. In this way, replay works synergistically with BTSP, which forms the synaptic connections that allows replay to occur after as little as a single exposure to novel information, helping to extend new learning without requiring additional exposure to environmental contingencies.

Beyond incorporating PS weighting into the theory and using BTSP as the key learning mechanism, we also extend the way the SR theory links the cognitive map to environmental features. The original SR theory links environmental states (places in an environment) to a simple scalar estimate of value based on reward, limiting its generality and adaptive flexibility^44^, since value of an external stimulus can vary with the animal’s needs and transient goals. For example, it would be adaptive to seek states associated with water when thirsty and to seek states associated with food when hungry. More generally, it would be a powerful extension of the SR theory to be able to use a SR to determine how to navigate toward or away from arbitrarily specified environmental features when they become goal relevant. To address this, we incorporate successor features (SFs) previously proposed in a machine learning context^45^ (another recent extension of the SR theory^46^ incorporates SFs to address issues different from those considered here). Unlike the SR, which predicts the *future occupancy of states*, SFs predict *the future occurrence of features* (Fig. 1b, bottom) that occur in these states. We propose that the PS-weighted SR is used to predict SFs in brain areas downstream from the hippocampus. As we shall see below, a simple error-correcting learning rule for learning feature predictions based on the PS-weighted SR speeds learning to predict salient features relative to others with lesser salience. The resulting feature predictions can then be assigned value based on the animal’s motivational state (or, more generally, its transient goals) during later behavioral episodes, allowing behavior to be guided toward places with current motivation-relevant properties.

Combining these components, we propose an extended integrative framework for the cognitive map, its neural implementation, and its consequences for rapid adaptation of neural activity and behavior (Fig. 1c-g). In this framework (Fig. 1c) the hippocampus integrates spatial input with PS signals to form a PS-weighted SR, which is then conveyed to downstream areas that compute SFs and combine them with motivational signals to estimate state values (Fig. 1d). To demonstrate how this representation can emerge during behavior, further strengthen during offline periods, and support rapid behavioral adaptation, our simulations employ three distinct phases: an online exploration phase, during which BTSP updates hippocampal connectivity (Fig. 1e); an offline replay phase, during which spontaneous replays propagate predictive structure and further enhance learning through synaptic plasticity (Fig. 1f); and a behavioral phase in which the learned predictive map supports flexible value computation and guides behavior (Fig. 1g).

## Results

### Formal statement of theory

Following the proposal of Dayan^8^, the SR for a starting state *s* and a target state *s*^*′*^ is the expected discounted future occupancy of *s*^*′*^ across time *t*, given *s*_0_ = *s*:

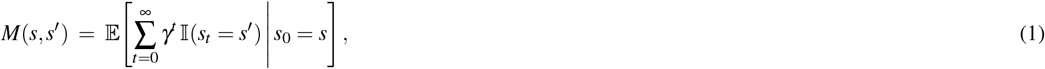

where *γ* ∈ [0, 1] is a discount factor. Thus *M*(*s, s*^*′*^) weights nearer states more heavily than distant ones. The Matrix *M* of current-state, future-state pairs summarizes the SR for a given environment.

The SR can be extended to characterize successor features, which correspond to discounted future predictions of environmental features (e.g., cues, rewards, or hazards)^45^. The SF estimate *ψ*(*s, f* ) for a feature *f* at start state *s* is defined as the expected discounted future occurrence of that feature:

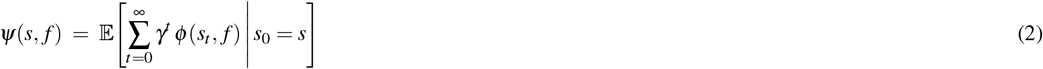

Which is simply the future occupancy of *s*^*′*^ times the estimated probability *ϕ* (*s*^*′*^, *f* ) of the feature’s occurrence at state *s*^*′*^:

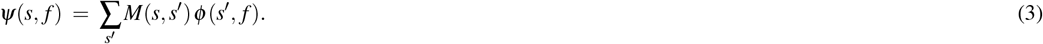

We next incorporate perceived salience (PS) into our framework^14,44^. The PS of state *s, ω*(*s*), is a positive quantity that depends on the sum, across features of a state, of the PS of each feature of the state, which in turn depends on the intensity, novelty, and motivational significance of each feature. We can then define the PS-weighted SR,

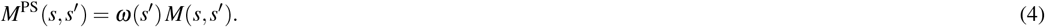

With this scaling, the SF representation *ψ*(*s, f* ) of feature *f* in state *s* can be computed by a linear mapping 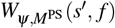 from *M*^PS^:

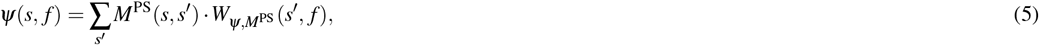

Note that this equation would compute the same successor feature probabilities as Equation 2 if the mapping weight 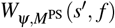 of PS-weighted state occupancy probabilities to feature predictions is set equal to *ϕ* (*s*^*′*^, *f* ) divided by *ω*(*s*^*′*^):

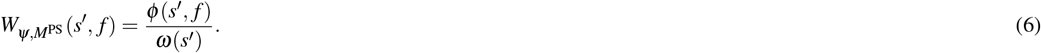

In our proposed implementation, the learning rule discussed below for estimating 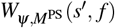 implicitly performs this division, while allowing predictions of features of salient states to be learned more quickly.

Finally, the value of state *s* at at some time *t* is defined as the sum of successor feature probabilities at *s* weighted by their value at the time of assessment

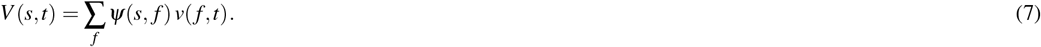

This allows the same predictive representation *ψ* to flexibly support different decisions, depending on the agent’s assessed value of different features when making decisions.

### Linking the theory to its neural implementation and behavior

We link these theoretical quantities to neural processes and behavior as follows:

- An estimate of the **PS-weighted SR**, *M*^PS^(*s, s*^*′*^), is acquired through BTSP in simulated neural populations representing hippocampal areas CA3 and CA1 during exploration and is further reinforced through replay-driven plasticity.
- **Perceived salience**, *ω*(*s*), thought to be computed outside of the hippocampus^14^, is computed as a simple function on the intensity, novelty, and motivational significance of features present at state *s* and is incorporated into the CA1 representation by modulating BTSP.
- The **mapping from PS-weighted SR to SF**, 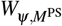, is instantiated by learned synaptic weights from the CA1 area to downstream feature-encoding populations. In our implementation, the learning rule is a simple error correcting learning rule^47^. As we discuss below (see Fig. 3), the up-weighting of the neural representation of the successor representation by PS speeds up the learning of features associated with highly salient states, contributing to rapid adaptation, while error correction prevents over-prediction.
- **State value**, *V* (*s*), is thought to be computed in extra-hippocampal circuits integrating feature predictions with feature-specific value signals *v*( *f* ), which we treat as scalar variables that can vary depending on the animal’s needs and goals at different times.
- The resulting *V* (*s*) guides **decision making**: when selecting actions, we simulate behavior by assuming animals preferentially transition to states with larger *V* (*s*).

In the following sections, we modeled neural activity and behavior after simulated exposure to environments paralleling those used in published experiments. We used distinct simulation paradigms for the online exploration, offline replay, and behavior phases of these simulations (Fig. 1e–g) as described more fully in *Methods*.

### Representations in CA3 and CA1 regions emerge rapidly through BTSP, supporting replay and prediction of successor features

We extended a spiking neural network modeling hippocampal areas CA3 and CA1^48^ and added circuitry to learn SF representations. The model includes a CA3-like population of 3200 simulated neurons with recurrent weights *W*_CA3_ and a CA1-like population of the same size with feedforward weights *W*_CA1_ from CA3 to CA1. As in previous work^48^, each CA3 neuron receives place-specific input in the form of a pre-specified Gaussian bump, with the centers of the bumps evenly covering a simulated linear track. On the other hand, CA1 neurons only received input via *W*_CA1_ from CA3. We added a downstream “feature-prediction” population of neuron-like units, each of which was assigned to correspond to a specific environmental feature, with weights *W*_FP_ projecting to it from CA1. We divided the linear treadmill into eight consecutive states (states 1–8), each of which was associated with a unique neutral “background” feature to distinguish spatial locations (Fig. 2a).

**Figure 2.**
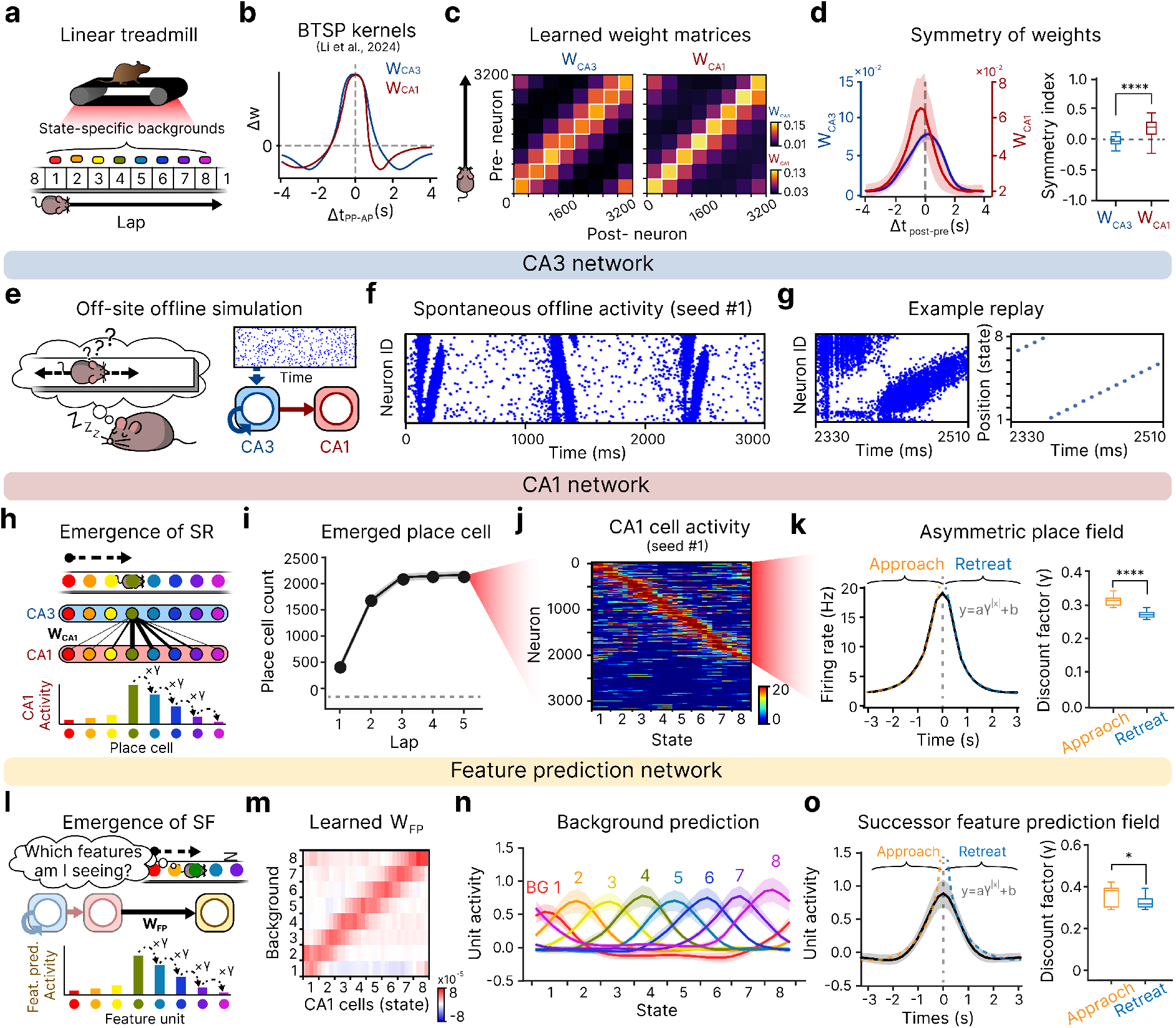
**a**, Linear treadmill simulation setup. **b**, BTSP weight update kernels for CA3 and CA1. **c**, Weight matrices after lap 5 with place cells sorted by field location, binned into 8 groups and averaged. Left: *W*_CA3_. Right: *W*_CA1_. **d**, Left: Averaged synaptic weights of postsynaptic neurons aligned by presynaptic index, for *W*_CA3_ (blue) and *W*_CA1_ (red). The curve is divided into weights from presynaptic neurons with smaller indices (“behind”) and larger indices (“ahead”). Scales for curves are chosen to aid visualization of differences in shape. Right: Symmetry index of *W*_CA3_ and *W*_CA1_, calculated by normalizing the area of behind vs. ahead weights. Paired t-test, *N*_*seed*_ = 10, \*\*\*\**P* = 4.58 *×* 10^−36^. **e–g**, Results from CA3. **e**, Offline simulation schematic. **f**, Example spontaneously generated offline activity. **g**, Left: Example replay activity. Right: Decoded position. **h–k**, Results from CA1. **h**, SR emerges in CA1 due to asymmetry of *W*_CA1_. **i**, Number of CA1 place cells formed after each lap. **j**, CA1 place fields after lap 5. **k**, Left: Average firing rate across place fields. Right: Fitted discount factors for approach vs. retreat trajectories relative to field centers. Paired t-test, *N*_*seed*_ = 10, \*\*\*\**P* = 7.02 *×* 10^−5^. **l–o**, Results from the feature prediction layer. **l**, Learning *W*_FP_ to predict SFs. **m**, *W*_FP_ after lap 5. **n**, Predictions for each background feature across positions after lap 5. **o**, Average predicted presence for the current state vs. non-current states. Paired t-test, *N*_*seed*_ = 10, \*\*\**P* = 8.13 *×* 10^−4^. All data are presented as mean*±*s.d.

We modeled BTSP-driven online learning through 5 contiguous laps of navigation— uniform movement through the assigned CA3 place field centers. During the online learning, *W*_CA3_ was trained using the BTSP learning rule with a symmetric time kernel^12,31^, whereas *W*_CA1_ was trained using an asymmetric kernel (Fig. 2b)^40^. *W*_FP_ was trained using an error correcting delta rule^47^ to predict the presence of the eight background features associated with the track states (Fig. 2l).

We first examined the learning of *W*_CA3_ and *W*_CA1_. As expected, the distinction in learning kernels resulted in different structures in these connections. To illustrate this, we separately identified CA3 and CA1 neurons by rank-ordering them according to the position at which each neuron exhibited its peak firing rate, from left to right along the track. Then, we plotted *W*_CA3_ and *W*_CA1_ after lap 5, arranging both pre-synaptic and post synaptic neurons by their IDs (Fig. 2c). The results showed that *W*_CA3_ encoded spatial relationships between place fields in an allocentric, movement direction-independent manner, resulting in a symmetric structure (Fig. 2d, blue). In contrast, *W*_CA1_ displayed a skewed distribution: connections from place cells representing positions behind the animal’s current location were more potentiated than those ahead (Fig. 2d, red).

Next, we examined the functional consequences that ensued from these distinct representations. We first showed that online learning through the symmetric BTSP rule enabled the CA3 network to spontaneously generate replay-like activity during subsequent offline periods. To test this, we simulated the offline phase by delivering a uniform, non-spatial input for three seconds to *W*_CA3_ after training (Fig. 2e). We simulated the resulting spontaneous activity using biophysically plausible parameters and an adaptive exponential integrate-and-fire neuron model (Supplementary Materials, Section 3)^48^.

During simulated replay, the CA3 neurons fired in sequence according to their place-field locations (Fig. 2f). These sequential events were best decoded as a linear traversal along the track and therefore qualify as replay events^49^ (Fig. 2g). The replay activity began to emerge after a single track traversal and was robust after 5 laps of exploration (few-shot learning)^38^.

Next, we showed that the asymmetric structure of *W*_CA1_ endows CA1 neurons with SR-like place fields (Fig. 2h) after 5 laps of learning from random initial weights. We classified a CA1 neuron as a place cell if its peak firing rate exceeded 15 Hz (75% of the model’s specified maximum of 20 Hz) at any position along the track (see *Methods* for details). We found that an average of 2,148 CA1 place cells had emerged (Fig. 2i). The resulting place fields were distributed across the environment (Fig. 2j). As expected, these place fields were asymmetric: neurons began firing earlier as the animal approached the field’s peak location and shut off more rapidly as it moved away (Fig. 2k, left). This was assessed by modeling the place cell activity during the “approach” and “retreat” from the center of the receptive field using the function *y* = *a × γ*^|*x*|^ + *b* (Fig. 2k, right), where the larger discount factor *γ* during approach compared to retreat reflects the expected asymmetry.

Finally, we demonstrated that the feature prediction network successfully learned to anticipate environmental features (Fig. 2l). As expected, by the end of the fifth lap, *W*_FP_ had become state-specific, with CA1 place cells preferentially projecting to the feature unit associated with their place field (Fig. 2m). Consequently, each background feature unit exhibited strongest prediction when the animal occupied its corresponding location (Fig. 2n), along with anticipatory activation similar to that seen in the CA1 place fields (Fig. 2o).

### Salient features are rapidly encoded and updated in CA1, enhancing their prediction in the feature prediction network

Next, we demonstrated that the SR encoded within the CA1 layer in our model reflects the introduction and relocation of reward^12,16,19^, one type of salient outcome, leading to rapid adaptation of place fields and outcome predictions. We introduced a food reward as a salient outcome after the five familiarization laps without reward described above (Fig. 2). The food was placed at state 4 for the next five laps (laps 6-10), and then relocated to state 8 for five additional laps (laps 11-15) (Fig. 3a). We treated the introduction of reward as affecting two key factors (Fig. 3b). First, the motivational importance of food was set to 3, compared with 1 for background features. This significantly increased the PS of the reward location, modulating the place field formation in CA1 (Fig. 3c). Second, the animal’s travel speed was halved when passing through the reward state, reflecting slowing during reward consumption^12,19^.

**Figure 3.**
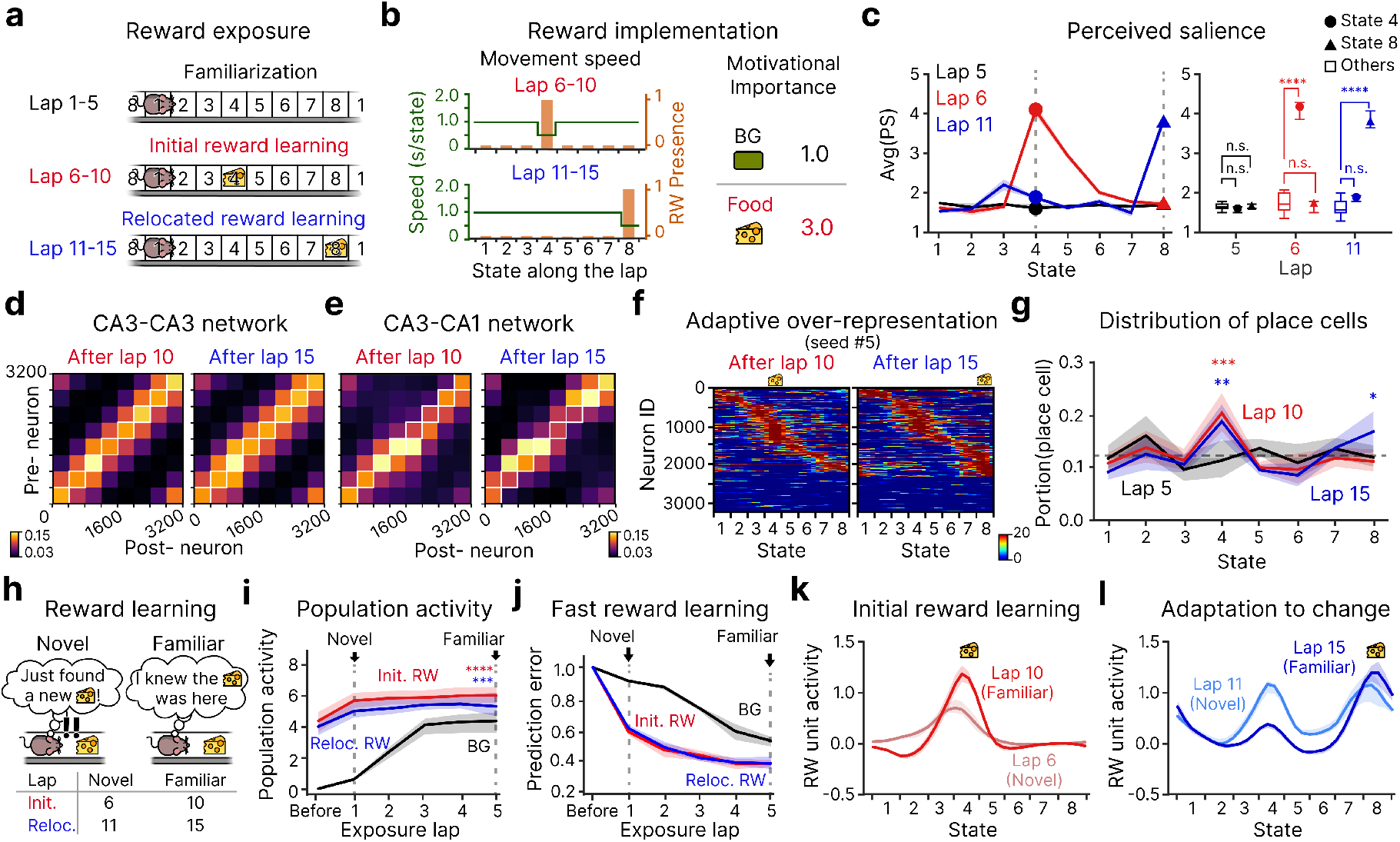
**a**, Simulation scheme for introducing a salient reward. **b**, Two factors modulated by reward introduction: reduced movement speed and increased motivational importance. **c**, Left: Average PS across states during laps 5, 6, and 11. The rewarded state exhibits elevated PS when the reward is present. Right: PS at states 4, 5, and all other states during laps 5, 6, and 11. Paired t-test, *N*_seed_ = 10, n.s. *P >* 0.05, \*\*\**P <* 0.005. **d**, *W*_CA3_ after lap 10 (initial reward introduction) and lap 15 (reward relocation). Values are averaged across 10 random seeds. **e**, *W*_CA1_ after lap 10 and lap 15, averaged across 10 random seeds. **f**, CA1 place cell activity after lap 10 and lap 15. Place fields cluster around reward locations and adapt following reward introduction and relocation. **g**, Proportion of CA1 place cells representing each state after lap 5, lap 10, and lap 15. The dashed line indicates the expected proportion (1/8) for a uniform distribution. One-sample one-tailed t-tests comparing mean *<* 1*/*8, *N*_seed_ = 10; \**P <* 0.05, \*\**P <* 0.01, \*\*\**P <* 0.005; unmarked comparisons are not significant with *P >* 0.05. **h**, Two cognitive states associated with reward perception. A reward is initially experienced as novel (left) and becomes familiar after repeated exposures at the same location (right). **i**, Development of CA1 population activity at background and reward-related states as the animal becomes familiar with environmental features. Population activity was computed as the mean firing rate across all CA1 neurons. The black line shows activity at state 4 from the initial network through lap 5, the red line shows activity at state 4 from lap 5 to lap 10 (reward present), and the blue line shows activity at state 8 from lap 10 to lap 15 (after reward relocation). Paired t-test, *N*_seed_ = 10, ****: CA1 population activity at state 4 at at lap 10 vs. CA1 population activity at lap 5, *P* = 5.70 *×* 10^−7^, ***: CA1 population activity at state 8 at at lap 15 vs. CA1 population activity at lap 5, *P* = 0.004. **j**, Prediction error for background features and the reward feature across laps. Background features require more laps to learn than the reward feature. **k**, Reward learning following initial introduction at state 4. **l**, Reward learning following relocation to state 8. All data are presented as mean *±* s.d.

In accordance with experimental findings and existing modeling work^12,31,40^ our model incorporates PS as a modulator of plateau potentials in CA1 and not in CA3. As expected, then, *W*_CA3_ was not meaningfully altered by reward relocation (Fig. 3d), while the predictive map encoded within *W*_CA1_ captured the PS information introduced by the reward. Indeed, *W*_CA1_ adapted rapidly to changes in reward location (Fig. 3e). After 5 laps with the reward in state 4, connections in *W*_CA1_ to CA1 neurons in locations near state 4 were strengthened, and after 5 laps with reward at state 8, connections *W*_CA1_ projecting to CA1 neurons representing this new reward location increased sharply. This adaptive reweighting arises from the modulation of BTSP in CA1 by perceived salience (Supplementary Fig. 3). As a result, the CA1 neurons were adaptively remapped to over-represent the vicinity of the reward (Fig. 3f and g), in accordance with experiments^15,16,31^. Notably, CA1 population activity at a rewarded state increased rapidly following a novel reward encounter (Fig. 3i), and remained approximately stable thereafter. This contrasts with the slower initial buildup at these and other sites when the have only non-salient background features.

We next considered changes in reward prediction, comparing predictions immediately following reward introduction or relocation (‘novel’, Fig. 3h, left) to predictions after repeated exposure (‘familiar’, Fig. 3h, right). Prediction error for the reward decreased much more rapidly than initial learning of background features (Fig. 3j). Furthermore, immediately after the reward was introduced at state 4, the network started to predict the reward at the reward location and to anticipate it at the preceding state (Novel; Fig. 3k). This prediction was gradually strengthened with repeated exposure to the reward at the same location (Familiar; Fig. 3k). After reward relocation to state 8, the feature-prediction layer immediately began to predict the reward at the new location (Novel; Fig. 3l). CA1 retained residual over-representation at the former reward site (Fig. 3f, right), and prediction at the old site decayed gradually with further experience (Familiar; Fig. 3l). This asymmetric adaptation reflects rapid acquisition of new salient associations^1^, alongside slower decay of memories about the previous location as supported by behavioral findings^50,51^.

### Learning to seek rewards relevant to current motivational state

We next explored how our framework supports using learned feature predictions to obtain rewards, highlighting use of these predictions to seek rewards relevant to an animal’s motivational state at the time when it is behaving. Using a setup similar to one used in an experiment demonstrating state-dependent reward seeking^52^, we trained a CA3-CA1 network with 4000 simulated neurons in each area on a two-dimensional T-maze with two distinct arms divided into 10 states. During the exploration phase, we simulated alternating visits to each arm, starting from the base of the stem (state 1), with odd-numbered laps ending at the end of the left arm (state 4), and even laps ending at the end of the right arm (state 10) (Fig. 4a). After six laps (three exposures to each arm), distinct reward features (food and water) with motivational significance of 3 were introduced at the arm ends, and the animal was exposed to each reward twice over the next 4 laps (7–10) (Fig. 4b).

**Figure 4.**
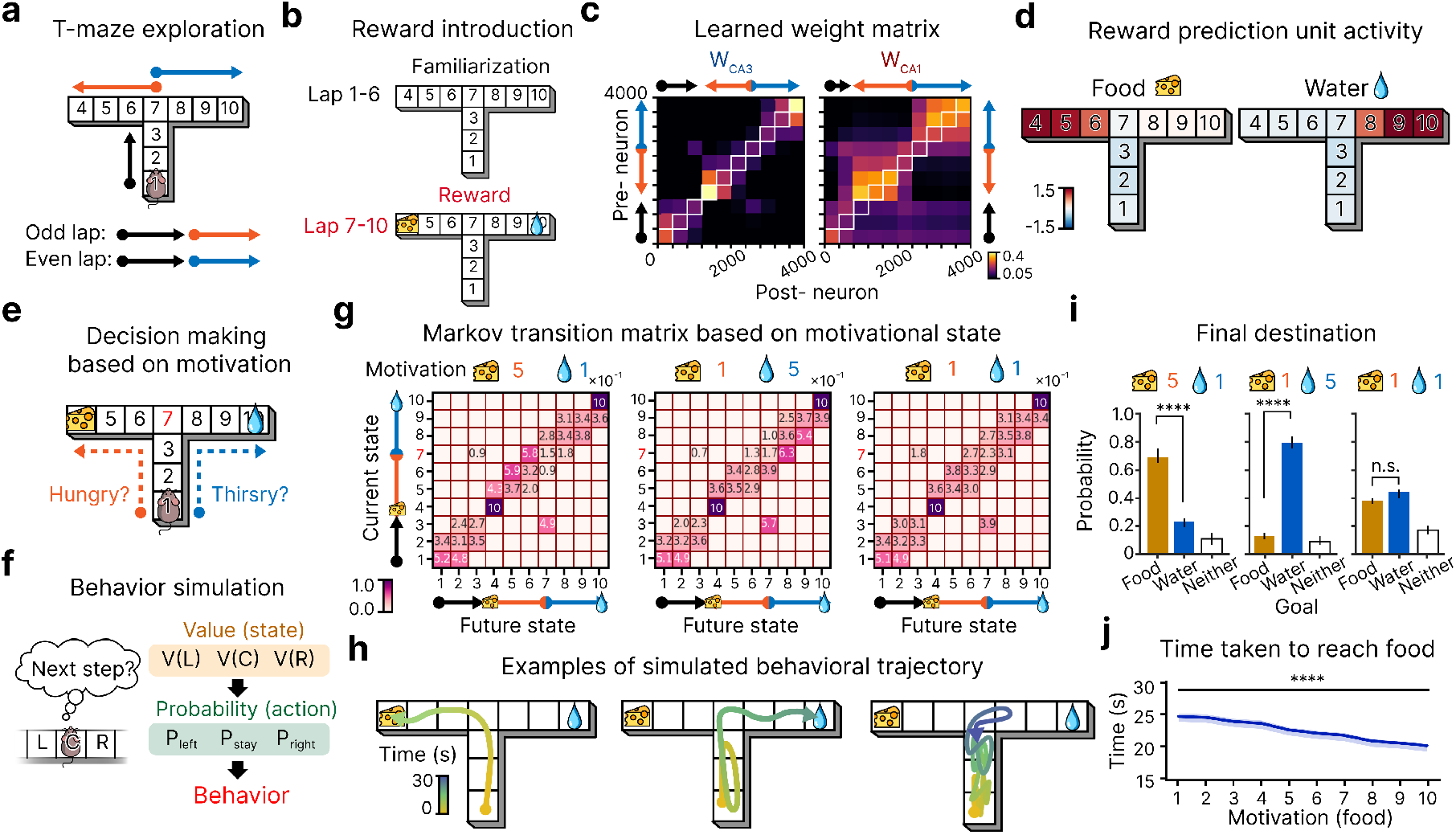
**a**, T-maze learning scheme. **b**, After 6 familiarization laps, two distinct rewards (food and water) are introduced at each arm for 4 laps. **c**, Resulting *W*_CA3_ and *W*_CA1_ after lap 10. **d**, Predictions for food and water across maze positions. **e**, Animal’s decision to pursue either arm of the maze based on its motivational state. **f**, Behavioral decision-making simulation scheme. The choice-making point (state 7) is marked red. **g**, Markov state transition matrices under three different motivational scenarios for food and water. Transition probabilities at state 7 shows most prominently the role of motivational state and therefore marked red. **h**, Example simulated trajectories for each scenario. **i**, Goal reached within 50 seconds of movement for each scenario. Paired t-test, *N*_*seed*_ = 10, \*\*\*\**P <* 0.0005, n.s. *P* = 0.98. Data are presented as mean*±*s.d. **j**, Time to reach the goal as a function of motivational strength. Mann-Kendall test, *N*_*seed*_ = 10, \*\*\*\**P* = 8.30 *×* 10^−5^, *Z* = −3.94. Data are presented as mean*±*s.e.m.

After lap 10, *W*_CA3_ encoded the structural layout of the T-maze (Fig. 4c, left), while *W*_CA1_ over-represented both arms, reflecting the presence of food and water (Fig. 4c, right). Correspondingly, the feature prediction layer strongly predicted each reward at its respective location (Fig. 4d).

We then simulated behavior based on the resulting learned map under different motivational needs for food and water (Fig. 4e). This simulation relied on value estimates *V* (*s*) for each maze state assigned in accordance with the formal theory (Equations 2-6). As described in detail in *Methods*, these estimates were obtained by projecting the center of each state through the Gaussian projection to produce an activity pattern in CA3 and propagating this through *W*_CA1_ and *W*_FP_ to obtain feature predictions *ψ*(*s, f* ) and assigning the final value based on the animal’s motivational state in accordance with Equation 6. These value estimates were used to assign next-state transition probabilities from each state to the set of state accessible from it using a softmax function (Fig. 4f). In each simulated behavioral episode, the animal started at the stem (state 1) and updated its location across a series of steps at a notional rate of one step per second based on the resulting Markov transition matrix. Each behavioral trial lasted up to 60 steps or until either goal state was reached.

We simulated three motivational conditions: (1) hungry (food need = 5, water need = 1), (2) thirsty (food need = 1, water need = 5), and (3) neutral (food need = 1, water need = 1). For each condition, we computed the Markov transition matrix containing the movement probabilities (Fig. 4g) as described above. The three matrices notably differed in the state transition probabilities from state 7 (row index highlighted in red) which is the choice point in the maze. When hungry, the animal was more likely to move toward the food arm (left), whereas when thirsty it preferred the water arm (right). Under neutral conditions, movements left, right, or back to the stem were roughly balanced. Accordingly, the animal predominantly reached the food when hungry, the water when thirsty, and distributed its choices more evenly under neutral conditions (Fig. 4h-i). Finally, we found that stronger motivational drives accelerated goal-directed behavior: as the motivational need for food increased from 1 to 10 (with water fixed at 1), the time required to reach the food decreased (Fig. 4j).

### Offline replay propagates perceived salience information to unexperienced paths, enhancing reward seeking from novel starting locations

We next investigated how learning during offline replay episodes can enhance the network’s prediction of salient outcomes and support behavior, including finding novel paths to reward, as observed in the seminal experiments of Tolman^2^. We used network weights learned through BTSP after exposure to food and water reward as described above (Fig. 4). Specifically, we hypothesized that replay activity propagates SF information across the maze, including to positions that were never visited in succession during online exploration^42^ (Fig. 5a). Accordingly, we simulated an offline phase corresponding to a post-reward pause at the food location by providing place input for state 4 to CA3 cells, then observed the resulting network activity. During this phase, replay spontaneously emerged (Fig. 5b, top). Strikingly, the replay events we observed propagated not only along previously experienced trajectories (left- and right-arm replays), but also exhibited a novel “shortcut replay” that had never been traversed, connecting the two arms of the maze (Fig. 5b).

**Figure 5.**
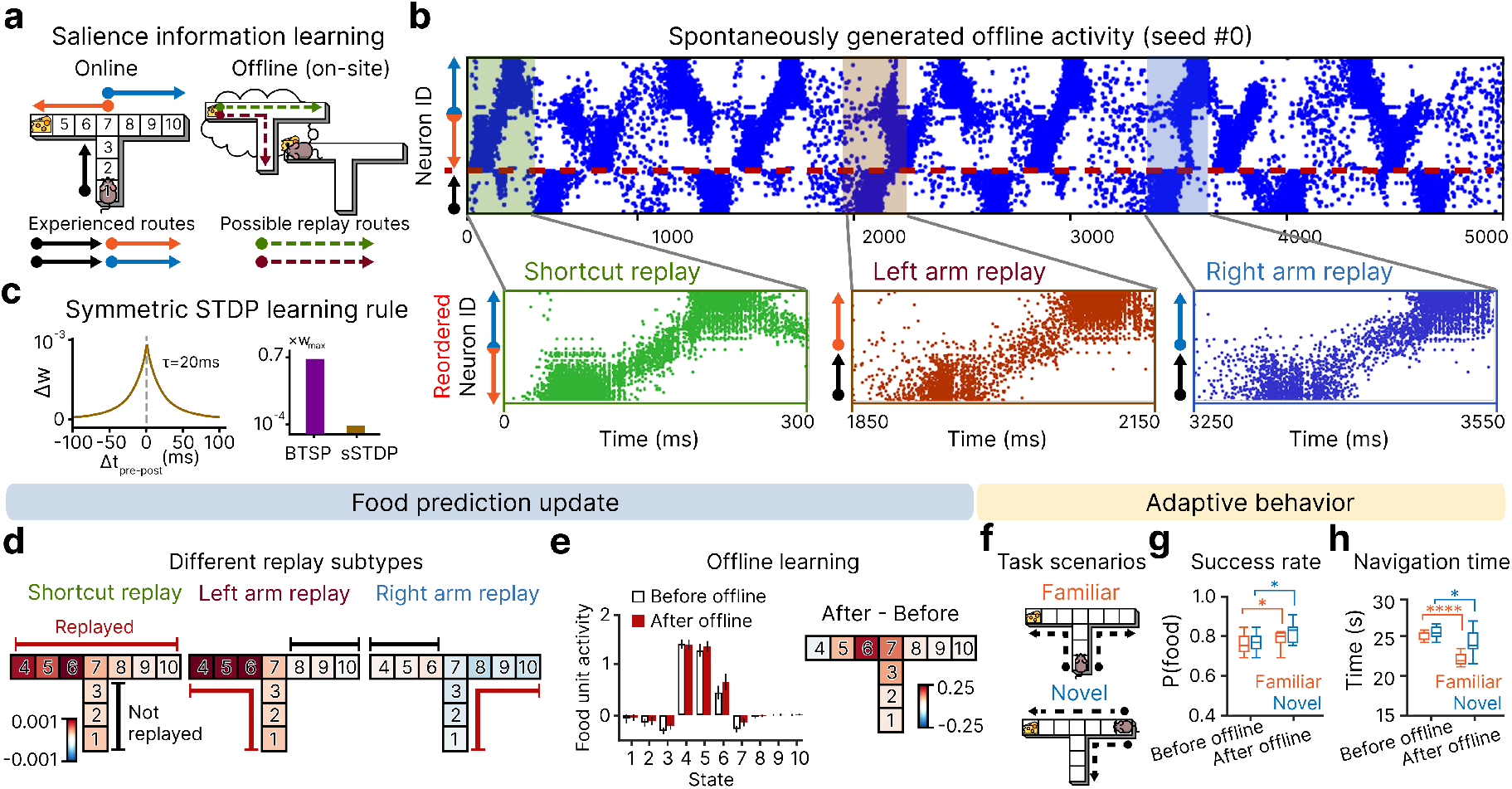
**a**, Propagation of PS information through online and offline learning. **b**, Spontaneously generated offline activity. Replay events are grouped into three categories based on their trajectories: shortcut, left-arm, and right-arm. For visualization, each expanded example shows only the neurons participating in the corresponding replay, reordered in the sequence in which they were activated along the replay trajectory. **c**, Left: sSTDP rule used in offline learning. Right: Largest possible weight update of BTSP and sSTDP rules. **d**, State-wise food prediction updates induced by a single replay event, shown separately for each replay subtype. Updates were computed as the difference between post-replay and pre-replay food predictions. **e**, State-wise food prediction updates induced by 5 seconds of offline learning. Left: Food unit activity at each state before (pre) and after (post) offline learning. Right: State-wise prediction updates computed as the difference between post- and pre-offline learning activity. **f**, Two behavioral simulation schemes. Left: Simulation starting from the same start position as during online exploration. Right: Simulation starting from a different start position. Both tasks aim to find food located at the same position as in the online exploration. **g**, Proportion of trials in which the animal reached the goal location within 50 seconds in familiar and novel scenarios, without and with offline learning. Paired t-test, *N*_*seed*_ = 10, Familiar \**P* = 0.03; Novel \**P* = 0.02. **h**, Time taken for the animal to reach the goal location in familiar and novel scenarios, without and with offline learning. Paired t-test, *N*_*seed*_ = 10, Familiar \*\*\*\**P* = 1.15 *×* 10^−5^; Novel \**P* = 0.04. All data are presented as mean*±*s.d.

We then examined how each replay type can contribute to propagating feature predictive information from the reward location to distant states. During replay, we used a temporally symmetric-spike-time dependent learning rule^48^ that adjusts synaptic weights after every spike with smaller magnitude and a narrower temporal window than BTSP (Fig. 5c).

To identify effects of distinct types of replay events, we determined the change in food prediction unit activity at each state, averaged over all replays of the same type observed during the 5-second replay period. Following shortcut replay events, food prediction increased across both arms of the maze. In particular, this update was most prominent at states proximal to the food location (states 4-7; Fig. 5d, left). Similarly, left-arm replay enhanced food prediction along the replayed trajectory (Fig. 5d, middle). In contrast, right-arm replay had minimal impact, as its trajectory does not include the food location and therefore does not directly strengthen synapses originating from food-related place cells (Fig. 5d, right). Finally, we updated *W*_CA1_ using the full spike trains recorded during the offline simulation containing multiple replay events of different subtypes. This offline update increased food prediction in reward-adjacent states, particularly those that had not previously strongly predicted food (Fig. 5e).

We then simulated two behavioral task scenarios. In the first, the animal started at the stem (state 1) with a motivational need of 5 (Fig. 5f, top), as during the online exploration phase (Fig. 4). Each trial ended when the animal reached either end of the maze (state 4 or state 10) or failed to reach a goal within 50 steps. In the second scenario, the animal started from a novel location (state 10) (Fig. 5f, bottom). Each trial ended when the animal reached either end of the maze (state 4 or state 1) or failed to reach a goal within 50 steps.

Across 500 trials per random seed, incorporating offline learning significantly increased the probability of reaching the food goal in both scenarios (Fig. 5g). Moreover, among successful trials, offline learning reduced the time required to reach the food in both cases (Fig. 5h), indicating that replay-driven updates not only improved success rates but also enhanced behavioral efficiency.

### The model learns aversive outcomes one-shot, predicts their presence through forward rollouts, and exhibits one-trial avoidance learning

We next asked whether the model could account for one trial avoidance learning, as observed in a study in which mice freely explored a linear track before encountering a footshock at the track end, after which they were promptly removed from the environment and placed in a rest box outside the linear track^1^. This differs from reward learning (Fig. 3–5) because animals rapidly move away from the shock, limiting the opportunity to associate the state and the outcome through prolonged exposure. Moreover, “on-site” replay is precluded by the animal’s rapid escape and immediate removal from the environment, although off-site replay may well have occurred in the rest box. In any case, animals exhibited one trial avoidance learning when placed back in the linear track after the rest period: none of the 4 animals entered the shock zone even once during a 10 minute period in the maze, though they had done so repeatedly in a session prior to shock presentation.

Our simulation approximated the design of this experiment. Simulated animals completed three laps of an eight-state environment without salient outcomes, followed by a fourth lap in which a foot shock occurred at state 8 (Fig. 6a). During the foot shock, we simulated movement at double the normal movement speed to align with the animal’s rapid escape responses. The foot shock was modeled as highly salient by assigning it a motivational importance of 10 (Fig. 6b). As a result, PS at state 8 increased sharply relative to prior familiarization laps (Fig. 6c), inducing over-representation of that location despite the brief dwell time (Fig. 6d).

**Figure 6.**
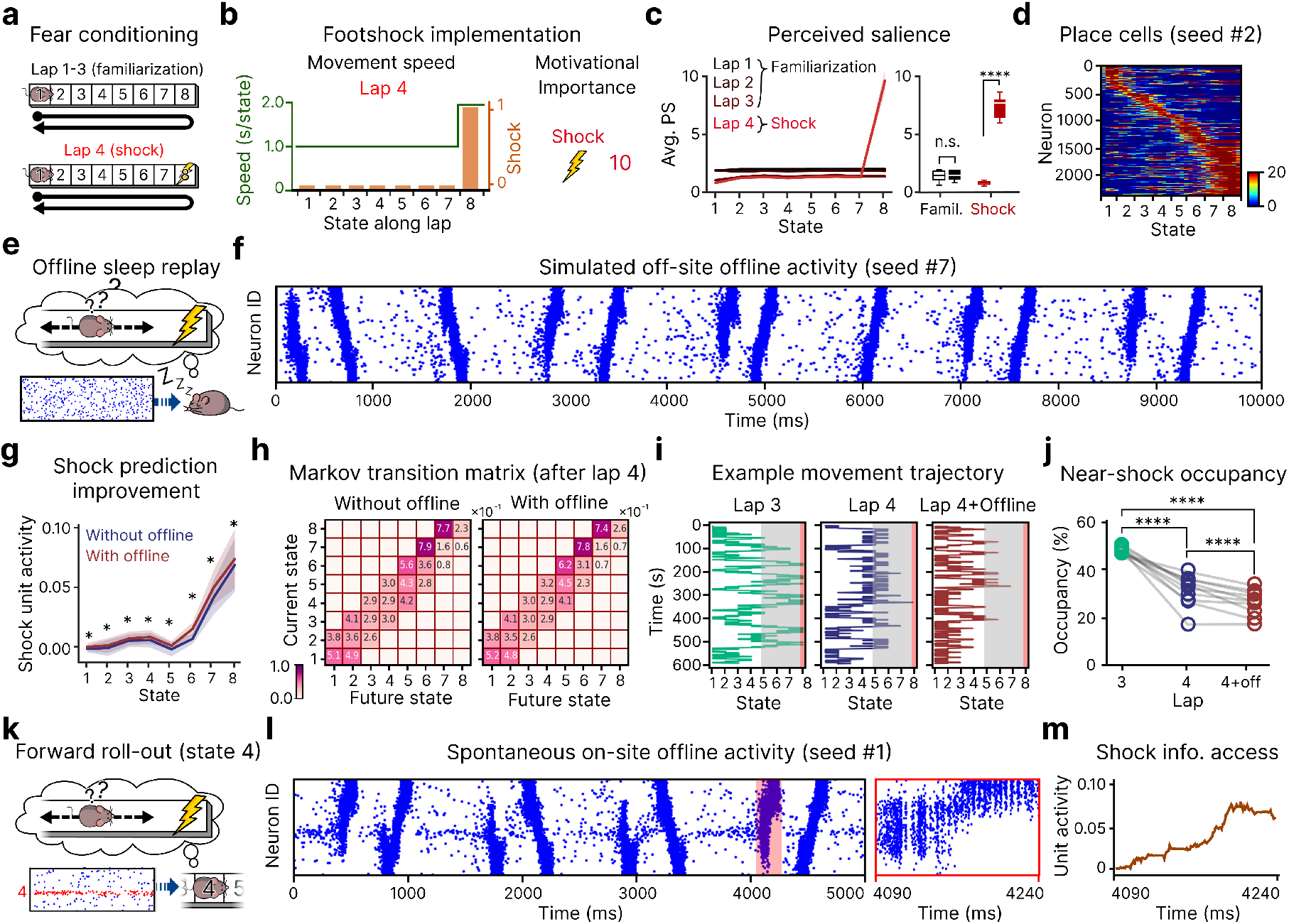
**a**, Aversive feature learning scheme. **b**, Two factors modulated by the introduction of aversive outcome: increased movement speed and the motivational importance of the aversive outcome. **c**, PS experienced at each state on lap 1 to 4. Independent t-test, PS of the non-salient states vs. PS of the salient state, *N*_*seed*_ = 10, n.s.*P* = 0.49, \*\*\*\**P* = 3.41 *×* 10^−64^. **d**, CA1 place cells clustered around the punished state, over-representing it. **e**, Offline simulation during sleep. **f**, Example spontaneously generated offline activity (10 s). **g**, Elevated anticipated footshock after the offline learning phase. Paired t-test, footshock prediction at each state without offline learning vs. with offline learning, *N*_*seed*_ = 10, \**P <* 0.05. **h**, Markov state transition matrices before and after the offline learning. **i**, Examples of simulated movement trajectories before footshock introduction (left), after footshock introduction without offline learning (middle), and after footshock introduction with offline learning (right). The red shaded region indicates the footshock state (state 8), and the grey shaded region indicates near-shock zone (states 5–8). **j**, Occupancy of the near-shock zone. Paired t-test, *N*_*seed*_ = 10, \*\*\*\**P <* 0.001. **k**, Forward roll-out at state 4, simulated by inputting state 4-specific input. **l**, Left: Example spontaneously generated offline activity (5 s). Right: Zoomed-in roll-out activity. **m**, Footshock prediction output from feature-prediction layer during the roll-out activity. All data are presented as mean*±*s.d.

We then simulated the subsequent offline phase after the animal was removed from the environment by providing random, low-intensity input to CA3 cells for 10 seconds (Fig. 6e). As a result, spontaneous replay-like activity emerged, spanning the entire track (Fig. 6f). We then used symmetric STDP to update *W*_CA3_ and *W*_CA1_ based on the resulting 10 seconds of simulated spiking activity^48^.

We found that incorporating offline learning extended the shock prediction field: replay enabled the shock to be predicted from locations farther away along the track (Fig. 6g). This change in prediction translated into altered behavior, simulated using the same method as in the T-maze simulation described above. Simulations of 600-seconds of free movement with and without inclusion of the offline replay based on the transition matrices shown in Fig. 6h revealed near-total avoidance of the shock zone even without replay. Offline learning further reduced the animal’s occupancy of the half of the track containing the footshock state (states 5–8; the near-shock zone) (Fig. 6i). We quantified this effect by measuring near-shock zone occupancy, which was significantly decreased when offline learning was included (Fig. 6j, right). Although the experimental study did not directly compare avoidance behavior with and without replay, our simulations highlight the possibility that the strong avoidance observed experimentally may depend in part on replay-mediated propagation of aversive predictions, as demonstrated in our simulations.

Another important finding from the same study was that, when the mice paused as they walked toward the shock zone after reintroduction to the environment, they exhibited replay events including forward-directed replays toward the shock zone^1^. These “forward roll-outs” may allow access to information about the foot shock without physically entering the foot shock zone. To test whether our model would also access this information after the single shock exposure and off-site replay described above, we simulated a second on-site replay phase with place input from the middle of the track (state 4) (Fig. 6k). The network spontaneously generated sequential place cell activity extending toward both ends of the track (Fig. 6l, left), including forward roll-out events that included the footshock state (Fig. 6l, right). Consistent with our hypothesis, internal footshock prediction increased during these roll-out events (Fig. 6m). These results show that our framework provides a basis for capturing the finding that aversive stimuli can affect neural responses after a single exposure to an aversive stimulus, highlighting distinct learning processes during behavior and offline periods that may synergistically contribute to adaptive behavior.

## Discussion

In our work, we have combined and extended the ideas and findings of many others to offer an integrative extension of the successor representation theory of the cognitive map and its role in value-based decision making, incorporating BTSP and hippocampal replay to address rapid neural and behavioral adaptation. Our work bridges abstract theory, neural representations, synaptic learning mechanisms, and neural and behavioral outcomes. We have built on the SR theory^8,9^ by incorporating successor features^45^: CA3 encodes a salience-neutral representation of environmental structure, CA1 integrates this with salience information to form a PS-weighted SR, and a downstream feature prediction layer computes SFs. The value of predicted features is assessed at decision time to guide behavior to address current needs and goals. The model is learned through BTSP^12,28,40,41^ during online exploration and further refined by symmetric STDP during offline replay, relying on circuit properties instantiated in previous models for replay propagation and learning^48^. It adapts swiftly when environmental contingencies change and captures a range of experimentally observed behaviors, including one-trial avoidance learning.

Our framework advances the seminal SR–based account of the cognitive map^8,9^ by separately considering PS^14^, feature prediction, and value assignment. These distinctions allow the model to capture the proposal that hippocampal networks over-represent salient locations, even when those locations are not inherently rewarding and do not lead to increased future visitation (Figs. 3 and 6). In addition, incorporating SFs, using them to represent environmental features including rewards, and separately assigning value to these features enables flexible integration of time-varying information such as motivational needs^52^. Because the model predicts features rather than states *per se*, it can adapt behavior based on motivational state without relearning the underlying environmental representation (Fig. 4).

Mechanistically, our work incorporates BTSP’s role in encoding distinct representations in hippocampal CA1 and CA3 areas^40^. In area CA1, our model extends the demonstration that temporally asymmetric BTSP can lead to emergence of SR-inspired place cells^28^ by incorporating PS weighting to capture the over-representation of rewarding locations^12,31^ and demonstrating how this can accelerate learning to predict salient environmental features. We further model how this over-representation may arise through different factors for rewarding versus aversive stimuli. In our simulations, positive outcomes were learned through moderate salience signals combined with prolonged occupancy (Fig. 3), whereas aversive outcomes were encoded through much stronger PS despite briefer dwell times (Fig. 6). Our model shows how these over-representations can accelerate learning of feature–location associations, as elevated activity at salient sites drives faster connection adjustments that support feature predictions (Fig. 3j). In CA3, on the other hand, BTSP learns a symmetric, value-neutral map of environmental structure through only a few laps of exploration, via temporally symmetric BTSP^40^.

Our model also highlights how offline replay can contribute to rapid learning by integrating several existing findings. Recent work suggests that symmetric recurrent weight profiles support spontaneous replay^41,48^, and that replay contributes to the consolidation of SR-like representations^21^. Integrating these results, our model shows that a value-neutral map encoded in CA3 connection weights can enable spontaneous replay after very limited experience (Fig. 2). Unlike online learning, replay sequences extend along unexplored trajectories^42,43^ (Fig. 5), enabling replay-driven inference in which information about salient features is propagated to states not actually visited in sequence, improving predictions. Beyond learning, our model incorporates a role for replay during behavior: simulated offline periods during immobility within exploration generate forward roll-outs similar to those observed experimentally^1^ (Fig. 6). These roll-outs transiently activate representations of non-current but salient states, enabling remote feature prediction and state-value estimation necessary for planning^49,53,54^.

Notably, our model highlights how BTSP offers an alternative to backpropagation of error signals though learned connection weights to recruit upstream neurons — in our case those in CA1 — to predict down downstream observations — in our case, successor features (Magee^26^ discusses related ideas, see Supplementary Materials Section 1.7). In our model, with BTSP, the signals causing CA1 neurons to change their incoming weights depend on a global PS signal, corresponding to the summed prediction error across all experienced features weighted by their intensity and motivational importance. This causes the model to produce upstream representations that support prediction without the transmission of error signals backward through connection weights or through precisely aligned return weight matrices, neither of which is thought to be biologically plausible^55^. The idea that a global signal might drive learning is not new. What is new is the large magnitude and extended temporal window of connection adjustments supported by BTSP, providing a mechanism that appears to allow an approximation of gradient-based learning to proceed far more efficiently than it could under previous proposals^56^. Indeed, this mechanism may accelerate initial learning relative to back-propagation, which requires forward weights to build up before they can carry error signals backward to drive upstream learning^57^. In sum, BTSP may endow a learning system with an implicit and highly efficient approximate error gradient propagation mechanism. This could go a long way toward explaining why biological brains learn far more efficiently than the gradient-propagation based learning systems used in today’s AI systems.

In spite of the progress our model represents, many issues and questions remain. We have made many simplifying assumptions to focus on building a conceptual bridge between levels and capturing rapid adaptation. Our simulations provide qualitative demonstrations rather than quantitative matches to details to provide a proof of principle. Future work should include additional pathways in the hippocampus and further details of the physiological implementation of some of the functional circuit properties of hippocampal and other circuits, to create more fully realized simulations of details of the underlying physiological mechanisms and a closer fit to some aspects of neural activity observed during behavioral and replay episodes (See Supplementary Materials Section 1 for further discussion).

Our work so far has focused on relatively simple learning settings. Future work should address more complex settings, including those requiring sensitivity to context. There is a very rich literature exploring how neural activity and behavior generalize from one context to another or differentiates between contexts, as explored in a recent extension of the SR framework^46^ and in work using a complementary computationally grounded framework that captures temporal context sensitivity^58,59^. We look forward to exploring whether our framework can be extended to capture the dynamics of representation learning and behavioral adaptation observed in these settings.

Broadening our focus, future work should address the learning efficiency gap^60^ between humans and today’s AI systems. As noted above, rapid online learning through BTSP and offline refinement through replay may offer alternatives to standard gradient-based updates, leading to a new understanding of how humans learn far more efficiently. It seems likely that BTSP contributes to the rapid formation of replayable mental trajectories that can be reactivated during offline periods in humans, as explored in recent investigations^61^. Insights from models employing BTSP and replay to better capture how these processes play out in human brains could contribute to the future development of more efficient learning methods in AI.

## Methods

### Neural network model

#### The hippocampal network

We modeled the hippocampal network as a two-layer spiking neural network, corresponding to the CA3 and CA1 subregions of the hippocampus, modifying and extending a simulation of exploration and replay in area CA3 from previous modeling work of Ecker et al.^48^ Consistent with experimental findings, the CA3 layer included recurrent connectivity (*W*_CA3_) and projected feedforward output to CA1 (*W*_CA1_). Synaptic weights in both pathways were constrained between 0 and 10, initialized from a Gaussian distribution (mean = 0.01, standard deviation = 10^−4^), and formed with a fixed connection probability of 0.1.

Both CA3 and CA1 layers contained the same number of excitatory neurons which were distributed across a small number of blocks we call ‘states’ tiling the simulated maze. Each state was represented by 400 neurons, with different numbers of states in the different environments we used as described below. To focus on learning processes within the hippocampus, we adopted simplifications of the inputs and plastic synapses in these brain areas. CA3 excitatory neurons were pre-specified to be place cells, with uniformly distributed place fields and spatial inputs reflecting the animal’s position during online exploration, and CA1 excitatory neurons did not receive direct spatial input; instead, their place-selective responses emerged through learned *W*_CA1_.

Dynamics of activity in this circuit differ between the online and offline simulation phases as described below.

#### Feature prediction layer

The feature prediction layer in our model aimed to provide a prediction of the physical presence of environmental features, including background, food, water, and shock. It comprised units representing the predicted presence (*ψ*) of each feature. These units received fully connected input from CA1 neurons via *W*_pred_ and allowed computation of a novelty signal (*N*) contributing to the modulation of the likelihood of plateau potentials in CA1 cells (See below). We interpret this network as a simplified representation of brain circuits involved in feature prediction and value computation, such as the nucleus accumbens, which forms a recurrent loop with CA1 and relays dopaminergic signals.

We now turn to our simulation of different phases of activity, again building on Ecker et al.^48^ Whereas these authors used STDP during behavior to set up subsequent offline replay and did not simulate further synaptic changes during replay, we use BTSP during online behavior and STDP during replay to capture the rapid adaptation that is the focus of our work.

### Online exploration

We simulated rodent’s maze learning task which is commonly studied in experimental literature. On each lap, we moved a notional simulated animal along a path through the environment while training the hippocampal and feature prediction networks.

#### Maze exploration scenarios

We simulated three different maze learning scenarios:

- **An animal unidirectionally navigating a linear-track treadmill**^**12**^: Each complete rotation of the belt corresponded to a 2.4 m lap. For convenience, the track was discretized into 8 states (state 1 to 8, from left to right), each 30 cm in length. In each lap, the animal began at the leftmost state (state 1), traversed to the rightmost state (state 8), reached the end of the belt, and was returned to the starting point. Formally,

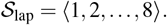

The next lap began immediately after the animal returned to state 1.
- **An animal navigating a T-maze**^**52**^: The maze consisted of a 0.9 m vertical stem (state 1 to 3, from bottom to top), connected at its midpoint to a 2.1 m horizontal arm (state 4 to 10, from left to right, with state 7 as the intersection). In each lap, the animal began at the base of the stem (state 1), moved up to the center of the horizontal arm (state 7), and then turned left in odd-numbered laps (ending at state 4) or right in even-numbered laps (ending at state 10). Formally,

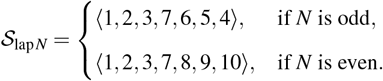
- **An animal round-tripping a linear maze**^**1**^: This scenario replicates an animal exploring a confined, 2.4 m segment of linear maze. The track was discretized into 8 states (state 1 to 8, from left to right), but unlike the linear treadmill, state 8 and state 1 are not connected. A single lap began with the animal at the left end (state 1), moving sequentially to the right end (state 8), and then returning back to the start (state 1). Formally,

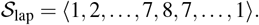

#### Environmental features

In our simulation, states were associated with distinct environmental features, *f* . Environmental features are accompanied with their physical intensity *I*_*f*_ as well as motivational importance *MI*_*f*_ . The presence and absence of the features at each of the designated states *s* of a given environment were encoded as a presence vector *P*(*s*) which remained constant across all positions within that state. Although real maze exploration environments contain many features, we simplified this to capture essential elements relevant to our work: background, food, water and shock.

- **Background:** Each state was assigned a state-specific background feature, which we thought of as the set of sensory cues in that state enabling the animal to distinguish the state from other ones. Since this feature is not attached to any affective value, we set the *MI* of this feature to be 1. When the state only has background but not other significant environmental features, the animal moves at the baseline speed, taking *T* = 1 second.
- **Food and water:** These features serves as a positive outcome in our simulation, and has *MI* of 3. When the animal was passing through the state with one of these features, it slowed to half the baseline speed, simulating an extended stay during reward consumption (*T* = 2 s).
- **Shock:** This feature serves as a negative outcome, with *MI* of 10. Once the animal enters a state where a shock occurs, it speeds up to twice the baseline speed for the shock state and subsequent states, reflecting an escape response (*T* = 0.5 s).

While stimuli can vary in intensity, for simplicity we set *I*_*f*_ to a constant unit value if feature *f* is present, so that *I*_*f*_ = *P*_*f*_ . Another component attached to environmental features is novelty, denoted as *N*_*f*_ (*t*). This quantity tracks the animal’s subjective surprise associated with feature *f*, and was defined as the difference between the predicted presence and actual presence of the feature. The novelty values were updated at each timestep based on the absolute prediction error computed by the feature prediction network, *ε*, described in the section below on training the feature prediction network.

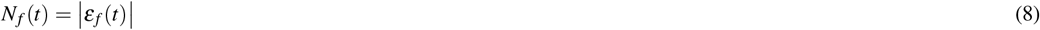

#### Computation of perceived salience

The perceived salience (PS) of an environmental feature *f*, denoted as *PS*_*f*_, represents the animal’s subjective evaluation of how noticeable or memorable *f* is. In our simulations, PS was modulated by three factors: motivational importance *MI*, intensity *I*, and the novelty *N* of the environmental feature. Specifically, the PS of feature *f, PS*_*f*_, was then computed as:

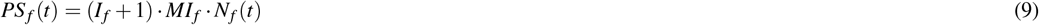

The added 1 in the expression (*I*_*f*_ + 1) is a fixed offset that allows the absence of a feature to affect learning. The PS of a state at time *t, PS*(*t*), was computed as the sum of PS across all environmental features.

#### Spike train generation and activity level estimation

CA3 place fields were uniformly distributed along the *x*- and *y*-axes, with each field centered at a distinct position. Place fields were circular with a radius of 30 cm.

Spiking activity in CA3 excitatory neurons arose from two sources: external spatial input from the entorhinal cortex, ℐ_place_, and recurrent input from other CA3 neurons, ℐ_recurrent_. *i*_place_ for a CA3 neuron depended on the animal’s Euclidean distance from the center of the neuron’s place field and followed a Gaussian profile. The maximum value of ℐ_place_ was fixed at 1 and the standard deviation of the Gaussian was chosen to be the radius of the field.

Recurrent input was computed as the sum of activities from all connected presynaptic neurons, weighted by *W*_CA3_. This summed input was then passed through a sigmoid nonlinearity to yield the final recurrent drive ℐ_recurrent,*i*_ for each neuron *i*. For a sigmoid function

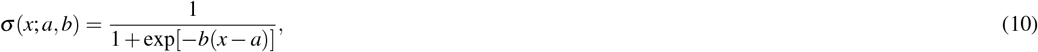

the recurrent input to CA3 neuron *i* was defined as

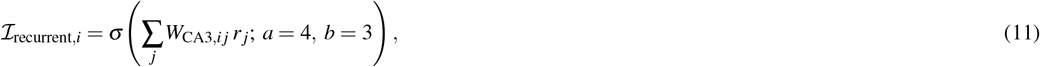

where ∑ _*j*_ *W*_CA3,*i j*_ *r* _*j*_ is the weighted presynaptic activity, and *a* and *b* are the sigmoid offset and gain parameters, respectively.

Finally, the total firing rate of neuron *i* was given by:

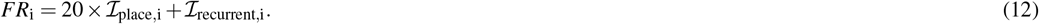

CA1 excitatory neurons were modeled similarly but did not receive direct spatial input. Their firing rates were determined exclusively by feedforward input from CA3, ℐ_feedforward_: the activity of CA3 presynaptic neurons was multiplied by *W*_CA1_ and passed through the same sigmoid nonlinearity (Eq. 10), with parameters *a* = 4 and *b* = 2. Formally, the feedforward drive to CA1 neuron *j* is:

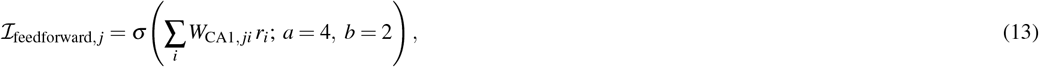

and the resulting firing rate is:

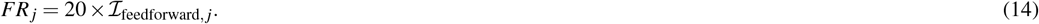

After calculating *FR*s of all neurons in both CA3 and CA1, we generated spike trains for each neuron using inhomogeneous poisson processes^28,48^. These spike trains were then filtered to remove spikes that violated an assumed 5 ms refractory period^48^.

To reduce noise in estimation of post-synaptic signals that arises from the small scale of our network, we introduced temporal smoothing: spike trains for both CA3 and CA1 were generated at every Δ*t*_spike_ interval, which was set to 100 ms. Neuronal activity 𝒜 was quantified as the actual average firing rate of each neuron, calculated by dividing the total number of generated spikes by Δ*t*_spike_. Note that this differs from the ideal *FR* values in the equations above, as it reflects post-processing by both the inhomogeneous poisson process and the refractory period filter^41,48^.

#### Training the hippocampal network using BTSP

Synaptic weight updates in the hippocampal network followed a BTSP rule, which depends on the interaction between presynaptic eligibility traces (ETs) and postsynaptic instructive signals (ISs)^19,31,40^. ETs were computed based on the spiking activity of presynaptic neurons, while ISs were driven by the postsynaptic neuron’s dendritic plateau potentials (PPs). When a presynaptic spike occurred, the post-synaptic neuron’s ET increased by a fixed amplitude, *A*_*ET*_, and then decayed exponentially with a time constant *τ*_ET_ = 1000 ms.

ISs were evoked by dendritic plateau potentials (PPs), modeled as stochastic events occurring in both CA3 and CA1 neurons. Each neuron *k* was assigned a plateau probability *p*_plateau,*k*_, updated every Δ*t*_spike_. At each 1 ms time step, a Bernoulli trial determined whether neuron *k* initiated a PP.

The plateau probability *p*_plateau,*k*_ was modulated by three factors: (1) the neuron’s activity 𝒜_*k*_ of the current Δ*t*_spike_ interval, (2) the perceived salience (*PS*) at the current time step, and (3) the mean activity of the entire layer, 𝒜_population_, which reflects layerwise inhibition and downregulates PP probability^19,31^ (Supplementary Fig. 1). Formally, the plateau probability for neuron *k* was:

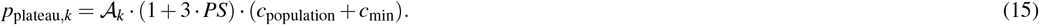

As CA3 layer did not receive a PS signal, *PS* was set to 0 and *c*_min_ = 0 for CA3 neurons. For CA1 neurons, *c*_min_ = 10^−5^.

The inhibitory scaling factor *c*_population_ was:

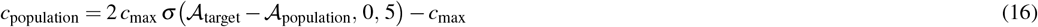

for 𝒜_population_ *<* 𝒜_target_, and

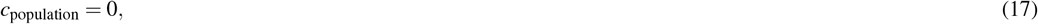

otherwise. We set *c*_max_ = 0.05 *×* (1 + 0.5 · *PS*) where *PS* was fixed to 0 for CA3 neurons.

When a PP occurred, the postsynaptic neuron’s instructive signal increased by *A*_IS_ and decayed exponentially with a time constant *τ*_IS_. For CA3 neurons, *A*_IS_ = 1.6 and *τ*_IS_ = 1000 ms, same as *τ*_ET_ which allows symmetric time kernel of BTSP. On the other hand, CA1 layer’s plateau potentials were modulated by P signal: *A*_IS_ = 0.4 *×* (0.25 · *PS*) and *τ*_IS_ = 500 ms for CA1 neurons, producing an asymmetric kernel^40^. BTSP-mediated synaptic updates occurred only during PP events. The BTSP update of the weights is a function of *ET × IS* as well as the initial value of the weight^31^. Initially weak weights undergo stronger potentiation even with smaller *ET × IS*, whereas the weights that are already strong enough are more likely to undergo depression even with stronger *ET* and *IS* signal. Formally,

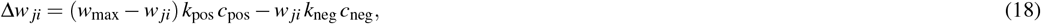

where

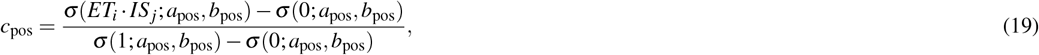

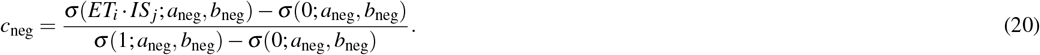

In our simulations, we set *k*_pos_ = 0.085, *k*_neg_ = 0.01, *a*_pos_ = 0.8, *b*_pos_ = 6, *a*_neg_ = 0.05, and *b*_neg_ = 44.44.

#### Training the feature prediction network using the delta rule

The feature prediction network was trained during the online exploration phase using the error-correcting delta rule^47^. Specifically, a synaptic weight *w*_*fi*_ connecting a CA1 neuron *i* and a feature unit *f*, was updated at each millisecond-level time step *t* as:

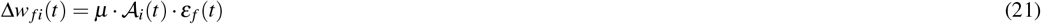

where *µ* is the learning rate, 𝒜_*i*_(*t*) is the activity of neuron *i* and *ε* _*f*_ (*t*) is the prediction error of feature *f* at time *t*. In our simulations, *µ* was set to 0.15 *×* 10^−7^ for the linear-track treadmill and linear maze scenarios, and 0.125 *×* 10^−7^ for the T-maze scenario. The prediction error was defined as the error between predicted presence (*ψ*) and the physical presence (*ϕ* ) for that feature:

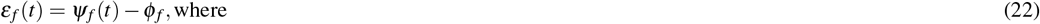

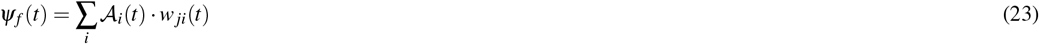

Finally, the novelty *N* associated with the feature at time step *t* is equal to the absolute value of the prediction error of the feature at that time step.

### Offline phase

The offline simulation framework was adapted from Ecker et al.^48^ Here, neurons were modeled using adaptive exponential integrate-and-fire dynamics, and synapses were modeled as conductances with biexponential kinetics (Supplementary Materials, Section 3.1 and 3.2). Hippocampal weights learned during online exploration (*W*_CA3_ and *W*_CA1_) were carried over into the offline phase, with additional connectivity to inhibitory interneurons introduced. Spontaneous activity emerging in the CA3 and CA1 layers during this phase was then used to update *W*_CA3_ and *W*_CA1_ according to a symmetric STDP rule.

#### Connectivity optimization

Additional connectivity was introduced and optimized using an evolutionary algorithm, including: entorhinal cortex input to CA3 excitatory cells (EC→Ext_CA3_); CA3 excitatory cell to CA3 inhibitory interneuron (Ext_CA3_→Inh_CA3_); interneuron–interneuron (Inh_CA3_→Inh_CA3_); and inhibitory interneuron to excitatory cell (Inh_CA3_→Ext_CA3_). The same connectivity types were established in CA1 (Ext_CA1_→Inh_CA1_, Inh_CA1_→Inh_CA1_, Inh_CA1_→Ext_CA1_).

The optimization algorithm minimized a loss function designed to match network-level physiological targets: maintaining realistic population firing rates in excitatory cells, suppressing gamma-band (30–100 Hz) oscillations, and enhancing ripple-band oscillations in excitatory cell populations (Supplementary Materials, Section 3.3).

During the offline phase, spontaneous activity was simulated in both CA3 and CA1, with low-rate entorhinal cortex input to CA3. *W*_CA3_ and *W*_CA1_ were updated using a symmetric STDP rule throughout this period.

#### Training the hippocampal network using symmetric STDP

During offline phase simulations with the optimized synaptic connectivity, we recorded the spiking activity of individual neurons in the CA3 and CA1 layers. These recorded spike trains were then used to update *W*_CA3_ and *W*_CA1_ according to a symmetric spike-timing–dependent plasticity (STDP) rule.

Symmetric STDP only involves potentiating the connections which connect neurons that fire together. This occurred in the second pass through of the recorded spike trains, using a time step of 1 msec. Precisely, the update rule we used is described as follows, with *A*_STDP_ = 1 *×* 10^−3^ and *τ*_STDP_ = 20 ms.

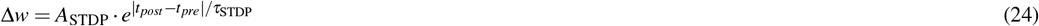

To simulate replay-based learning after exposure to food rewards in the T-maze (Fig. 5d), we isolated spiking activity within the temporal window corresponding to a single replay event. The learning effect was quantified by comparing the network’s predictions before and after the weight update induced by each replay event (Supplementary Materials, Section 3.4). Similarly, the learning effect after the full 5-second offline simulation was also simulated and analyzed using the entire spiking activity (5e).

To simulate replay-based learning after exposure to shock in a linear track (Fig 6g), weight updates were based on all of the spiking activity across the 10-second offline and off-site replay episode.

#### Estimating real-time feature unit activity during roll-out events

To estimate real-time feature unit activity in the feature prediction layer during offline replay (Fig. 6m), we used spike trains recorded during offline simulations. For each detected roll-out event, we computed CA1 firing rates using spikes within a 100 ms window preceding each time point, starting from the event’s onset time. This procedure was repeated at 1 ms resolution throughout the duration of the roll-out event. Then, these CA1 firing rates were multiplied by the feature prediction weight matrix *W*_FP_ learned during online training to estimate feature unit activity at every time point.

### Computation of successor feature activations

Here we describe how we determined the network’s representations of successor features Ψ _*f*_(*s*) for a given feature *f* at any location *s* in the environment based on the knowledge in connection weights before or after exploration and/or replay. These quantities are shown directly for 32 evenly spaced positions *s* along the linear track used in simulations shown in Figs. 2n and 3k,l and were also used as the basis for choosing actions in our behavioral simulations shown in Figs. 4 to 6.

We first computed CA3 neuronal activity associated with position *s*. A position- and input-dependent firing rate for each CA3 neuron was first determined by using its specified spatial tuning function relative to the current position. This was then scaled by a factor of 0.45 to approximate the influence of recurrent input through *W*_CA3_. CA1 neuronal activity was subsequently computed in the same manner as during the online exploration phase. Specifically, CA3 activity was forward-propagated through the feedforward weights *W*_CA1_ and passed through a sigmoid nonlinearity to obtain CA1 firing rates. Finally, feature-prediction unit activity was computed by linearly projecting CA1 activity through *W*_pred_, giving estimates of predicted feature activation Ψ _*f*_(*s*).

### Behavioral simulation

We conducted behavioral simulations using the information encoded in the network’s three weight matrices *W*_CA3_, *W*_CA1_, and *W*_FP_ network to examine the effects of learning on behavior. This phase corresponds to the exploitation stage in reinforcement learning, during which the agent uses previously acquired knowledge to make choices. No weight updating occurred during these simulations. We simulated two behavioral decision-making scenarios: one involving a positive outcome in a T-maze and another involving an aversive outcome in a linear track. In both cases, we treated the behaving animal as standing at the center of one of the discrete states in the relevant environment, and as computing an estimate of *V* (*s*) for the current state and for each state immediately adjacent to it in the environment. We used these *v*(*s*) values to determine next state transition probabilities, and then sampled from these probabilities to transition to the next state.

To estimate *V* (*s*), we first computed neuronal and feature-unit activity from the learned weights as described in the section above. The value of a given state *s*, denoted *V* (*s*), was computed as:

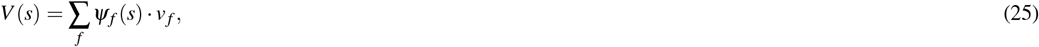

where *ψ*_*f*_ (*s*) represents the predicted presence of feature *f* at state *s*, and *v* _*f*_ denotes the subjective value of that feature. The feature values *v* _*f*_ were dynamically modulated by the animal’s transient motivational state. Given these computed state values, transition probabilities from each state to each other accessible state were computed using a softmax function. Specifically, given the vector of state values **V** = {*V*_1_,*V*_2_, …,*V*_*N*_} for the set of accessible states, transition probabilities were computed as follows:

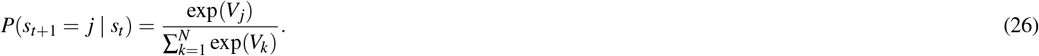

The simulated next state was determined by sampling from these probabilities.

#### Decision-making scenarios

- **T-maze with two goal locations and distinct rewards^52^**: Each trial began with the animal positioned at the stem of the maze (state 1) and ended when it reached either goal state (state 4 or 10). Food was located at the left end (state 4) and water at the right end (state 10), and the animal’s motivational needs for these two rewards were systematically varied. To test the network’s adaptability to different starting conditions, additional simulations were conducted in which the animal started from the right end (state 10) and traversed either toward the left end (state 4) or the stem (state 1), with food placed at state 4. In all cases, the animal was allowed a maximum of 50 steps per trial. Trials were terminated if the animal failed to reach a goal within this limit, and such cases were labeled as “failure to find the goal.”
- **Linear track with an aversive outcome^1^**: Each trial began with the animal positioned at the left end of the track (state 1) and allowed to move freely for 600 steps. A shock stimulus was positioned at the right end of the track (state 8).

### Statistical analysis

All statistical analyses were performed using custom Python code. Unless otherwise stated, data points represent independent simulation runs initialized with different random seeds (*N*_seed_ indicated in figure captions).

For comparisons between two conditions, we used paired t-tests as appropriate. One-sample t-tests were used when comparing a condition against a theoretical baseline.

Error bars represent mean *±* standard deviation (s.d.) unless otherwise noted. Exact *P* values and statistical tests used are reported in the corresponding figure captions.

## Acknowledgement

We thank A. Milstein (Rutgers University) and J. Magee (Howard Hughes Medical Institute) for valuable feedback on our implementation of BTSP and for their insights regarding the biological plausibility of our results. We also thank S. Jones, S. Grant, and J. Han (Stanford University) for helpful discussions during the early stages of the project. S.C. acknowledges support from the Emergent Cognitive Functions Fund, Department of Psychology, Stanford University.

