## Supplementary Materials for "Capturing rapid learning in an extended successor representation theory of the cognitive map"

### 1 Abstractions in our implementations of neural architectures and processes

Our work aims to bridge between abstract theory of the cognitive map and the neural processes that implement it in the brain, with a focus of capturing the rapid, adaptive changes observed in neural activity and behavior. To achieve this, we have intentionally abstracted away many anatomical, physiological, and circuit-level details of the brain, as well as properties of the experimental paradigms and the observed behavior. Incorporating additional experimental details to our model would produce better alignment with empirical data without altering its core conclusions. Below, we outline the main abstractions made when mapping biological circuits onto our model and discuss how including these elements in future work could further strengthen alignment with data.

#### 1.1 Input to CA3 area

To begin with, our model does not simulate the emergence of place cells in CA3; instead, we provide input to these cells that effectively assigns them to specific locations. In rodents, exposure to a novel environment rapidly induces CA3 neurons to develop distinct spatial and cue-related firing fields [24, 10]. This process likely depends on specific upstream inputs that are omitted from our model.

One major omission is input from the dentate gyrus (DG). CA3 receives sparse projections from DG via the mossy fibers which exert a very strong excitatory influence on the CA3 neurons they synapse on [1]. These have long been thought to play an important role in selecting neurons in CA3 to respond to the inputs they receive from other areas [36], and it has been suggested that they could contribute to inducing the dendritic plateau potentials necessary for BTSP to induce place fields and cue-modulated responses in CA3 [27]. Incorporating a DG granule cell layer could therefore provide a potential mechanism for determining which CA3 neurons become active place or cue cells.

Other omissions are grid cell and head direction input from the entorhinal cortex (EC). Grid cells provide periodic spatial basis functions, and combinations of these inputs are sufficient to generate single-peaked place fields [34, 43, 13]. Head direction input from the EC would allow CA3 place cells to exhibit directional selectivity, as is typically seen in place cells in linear tracks. Including EC grid and head direction input, together with DG drive, would likely provide a more realistic account of how CA3 place fields are formed during exploration.

Beyond spatial information, CA3 also receives environmental feature–related signals including their novelty and affective salience. These may arise either directly from the basolateral amygdala [51, 25] or indirectly through projections from CA1 and EC [22]. Such pathways could enable CA3 neurons to develop feature-related receptive fields alongside spatial ones.

Taken together, DG input, EC input, and other modulatory signals may support a richer cognitive map within CA3, allowing neurons to form multiple fields (both spatial and feature-based). This, in turn, could allow CA3 populations to represent spatial positions and features in partially distinct subpopulations, with recurrent synaptic strengths reflecting learned place–feature associations.

A likely downstream consequence of the place–feature associations in CA3 is their modulation on replay content. Because CA3 is a primary generator of hippocampal replay and sharp-wave ripples [6, 40, 50], combining spatial and feature-related inputs would be expected to bias replay toward trajectories involving behaviorally significant locations [49, 37]. One potential mechanism for the replay content modulation involves residual dopaminergic signals that persist during brief rest periods when the animal remains in the environment. Such signals have been shown to enhance excitability of neurons associated with salient states [33], increasing their likelihood of participating in replay.

#### 1.2 Input to CA1 area

As with CA3, a major omission in our CA1 implementation is the set of inputs it receives from the EC. Among these, head-direction signals are particularly relevant. Head-direction cells were first identified in the postsubiculum [46] and are known to encode the animal's facing direction independently of its spatial position. These head-direction inputs are thought to help distinguish locations that are visually similar but experienced from different orientations [21], and contribute to the directional tuning of hippocampal place cells. This directionality has been proposed to play an important role in shaping replay dynamics, particularly in distinguishing forward and reverse replay. In most of our simulations, the animal's movement during the online phase is constrained to a single direction along the track, and we do not model head-direction inputs or directional tuning of place cells. Incorporating head-direction signals and directional place coding in future models may provide a more complete account of how movement direction influences both spatial representation and replay structure.

CA1 integrates position and reward-related information from medial and lateral entorhinal regions, which convey both spatial structure and non-spatial feature cues [20, 47]. Together with the environmental-feature–specific inputs described above, these signals may contribute to the development of ”reward cells” in CA1 that fire selectively at the reward location, which firing fields shift to the new reward position when the reward is relocated [14].

We further propose that the increasing specialization of CA1 neurons could contribute to population-level orthogonalization based on task structure [45]. In this view, CA1 ensembles do more than representing spatial environmental structure. Rather, they progressively differentiate their activity patterns according to task-relevant cues, such as contextual cues that signal the current behavioral contingency [45]. Such task-dependent orthogonalization may reduce interference across task states that share similar spatial structure but differ in behavioral demands.

#### 1.3 Regulation of activity during online learning

Another area for refinement in our model is the fuller inclusion of physiological mechanisms that regulate neural activity, dendritic plateau potentials, and the resulting formation and activation of place cells in CA3 and CA1. Below, we highlight aspects of our simulations that could more closely align with neural data by including such mechanisms.

First, experimental studies show that place cells in reward zones tend to have narrower receptive fields than those in non-reward zones [4, 18]. In our simulations, by contrast, the place fields formed around these zones tend to have broader fields than those in non-reward zones (Fig. 3f). Mechanisms that more strongly regulate how many CA1 neurons can be active at the same timepoint might contribute to better capturing this detail of the experimental observations. We present a preliminary simulation consistent with this idea in Supplementary fig. 4.

A second point concerns the temporal symmetry of the BTSP weight-update kernel. Whereas our model treats instructive signals and eligibility traces in CA3 as having the same time constant, leading to temporally symmetric weight updates as observed in the first report of BTSP in area CA3 [27], recent investigation of BTSP in hippocampal area CA3 suggests that weight updates can be asymmetric during initial learning in a novel environment [29], consistent with the idea that the instructive signals underlying BTSP in CA3 can be shorter than synaptic eligibility traces.

#### 1.4 Regulation of activity during offline learning

A further simplification in our model are made in the offline simulations. In our simulations, offline activity was generated using an adaptive leaky-integrate-and-fire model, which incorporates firing-rate adaptation such that neurons active immediately before an offline epoch become less likely to fire. This mechanism has been proposed to influence replay directionality by biasing propagation away from recently active neurons [11, 3].

However, in our implementation, the firing activity from the end of the online exploration phase was not carried over into the subsequent offline simulation. Incorporating this continuity may help explain a broader range of experimental findings regarding replay content. For example, Wu et al. showed that when an animal pauses along the way to a salient aversive location, replay preferentially propagates forward toward the shock site [49]. If recent neural activity were inherited across phases, neurons representing positions just behind the animal (ones farther from the shock) would be less likely to participate in replay due to recent firing-induced adaptation, whereas neurons coding positions ahead (including the shock zone) would remain more excitable. This carryover mechanism may therefore help account for preferential forward replay toward salient outcomes during on-site pauses.

A related point is about the overall frequency and sparsity of replay events during offline periods. We found large differences in the frequency of replay events and the amount of activity within these events in our replay simulations (compare Figs. 2f and 5b). While the features of natural replay events certainly differ from setting to setting, these differences suggest to us that more refined regulation of neural activity within replay events could improve biological realisticity. Relatedly, replay events in our simulations are less sparse in our simulations than those observed in neural recordings. Regulation of these replays through inhibitory neural circuitry could potentially achieve more realistic sparsity.

Finally, there is evidence that the establishment of replay sequences depends not only on the formation of spatially- and temporally-adjacent place fields but also on fine-grained neural synchronization mediated by theta phase precession during the exploration episodes when the place fields themselves are being formed [28]. Our current simulations, though they include theta-phase modulation of neural activity, rely on a temporal smoothing over 100 msec time windows that tend to obscure this modulation. Our simulations are able to establish temporally-structured replay in spite of this, perhaps because we provide CA3 cells with strong spatial dependence due to the spatially aligned inputs, as described in *Methods*. Future work relaxing this pre-specified dependence would likely reveal the importance of spatial structuring through theta-phase precession. Future work should explore whether BTSP is a sufficient learning rule to exploit this structuring, or whether a spike-time dependent learning rule is required, as proposed in ref. [28]. Regardless of this, our simulation still captures the proposal that BTSP creates the necessary place fields needed as a crucial part of establishing replay.

#### 1.5 Roles of replay to support learning

Our work shows that both the content and frequency of replay can be modulated based on the property of the input signal (Figs. 2 and 6; Supplementary fig. 5). We further demonstrated that replay, depending on its trajectory, can selectively propagate salience information to the states it traverses (Fig. 5). Building on these results, future work could examine how replay content is regulated according to the animal's cognitive demands. For example, some studies suggest that replay prioritizes newly acquired salient memories [2, 49], contributing to their consolidation. Other work indicates that replay can preferentially sample trajectories that have not been recently experienced [19, 16, 7], potentially reinstating them to preserve older memories. Balancing these two seemingly competing functions—strengthening new salient information while maintaining previously learned material—is thought to be critical for preventing catastrophic interference during ongoing learning [32, 39, 48, 8]. How the hippocampal replay negotiates this balance, and how it is elicited to do so, remains an open question with contrasting theoretical perspectives. One perspective proposes that replays optimally address animal's needs and goals while planning [31], while another suggests that patterns of replay can be understood as reflections of the interplay of mechanistic processes that are together sufficient to explain the kinds and frequencies of replay events that are observed [52]. Future work more fully incorporating goal-related neural activity into replay dynamics could bridge these disparate perspectives.

#### 1.6 Feature prediction, value estimation, and computation of perceived salience

Outside of the hippocampus, our model simplifies the downstream structures involved in feature prediction in a single "feature prediction layer," and uses a simple mathematical expressions over variables represented in this layer to compute value and perceived salience. Although multiple interconnected regions contribute to these computations biologically, we do not implement the detailed mechanisms of these downstream circuits because our primary focus is the hippocampal learning mechanism.

In reality, several cortical and subcortical structures likely participate in these processes. One key area is the nucleus accumbens (NAc). Prior studies suggest that NAc encodes predictions about upcoming cues and outcomes and integrates motivationally relevant information for reinforcement learning [42, 17]. Importantly, NAc is reciprocally connected with the hippocampus: it receives dense glutamatergic projections from CA1 conveying spatial context and salient event information [38], and it can in turn modulate hippocampal processing through feedback pathways. Within our proposed framework, projections from CA1 to NAc could transmit salience-weighted spatial and feature signals, enabling NAc circuits to compute successor-feature predictions. Conversely, NAc could send novelty or prediction error signals back to CA1, helping to update the perceived salience-weighted successor representation.

Beyond NAc, multiple different brain subregions, such as the orbitofrontal cortex (OFC), ventral pallidum, and ventral tegmental area (VTA), have been implicated in computing or relaying feature-specific predictions, encoding state–space structure, and providing value- or novelty-related teaching signals [41, 12, 5]. These regions could work with hippocampus–NAc interactions to support flexible feature prediction and value computation. In summary, while our model abstracts these diverse downstream systems into a single feature-prediction module, future work incorporating their specific circuit dynamics may yield a more complete mechanistic account of how hippocampal representations guide adaptive behavior.

#### 1.7 Conceptions of the role of the inputs initiating plateau potentials in CA1

Here we briefly compare our conception of the role of input from outside the hippocampus in driving plateau potentials in CA1 with the conception offered in a recent review of BTSP by Magee [30]. In our conception, this input (implemented as the perceived salience signal in our model) is a weighted sum of novelty signals, each represented as the unsigned magnitude of prediction error of an environmental feature, a signal that is proposed to be computed by dopamine neurons in nucleus accumbens [23]. As can be seen in Equation 9 in *Methods*, this signal goes to 0 when environmental features are all predicted perfectly, although our model includes an offset from 0 to allow some plateaus to form even when there's no prediction error (Equation 15). This signal can be seen as informing CA1 that new place cells need to be recruited by the induction of plateau potentials; the recruitment or a particular cell is affected by its activation (depolarization) based on synaptic inputs from CA3 and potentially EC as well, so that cells with incipient tendencies to respond are more likely to have plateaus. Magee also describes the input driving plateau potentials as a global signal and ascribes a similar role to existing connections from CA3 to CA1 in selecting which CA1 cells will be recruited [30]. The difference in the proposals lies in our treatment of the input from outside the hippocampus as a salience weighted signal that something surprising has occurred, whereas in Magee's formulation this signal is described as a target signal that is modulated by importance (e.g. reward) without being modulated by prediction error, citing evidence that this signal does not decrease even when plateau probability has leveled off and the animals behavior indicates learned anticipation of reward [18]. While the evidence cited indicates that the EC input does not change, other work suggests that dopamine input from other brain areas influences plasticity in CA1 [26, 33]. This input may combine with the EC input signal to influence the probability of plateau potentials and/or the extent of plastic changes that are made when plateaus occur. In any case, both Magee's and our proposals share the view that the EC input is relatively global rather than specifically targeted to individual CA1 neurons, and both offer an alternative to the view that error signals need to target specific upstream neurons in selecting them to play a role in downstream predictions, as we discuss in our *Discussion* section.

### 2 Pseudo code description of the online learning process using BTSP

The complete online learning process is described as Algorithm 1 below.

**Algorithm 1:** Training during online environment exploration

**Initialize:**

    `# Initialize weights and signals.`
    $W_{\mathrm{CA3}} \sim \mathcal{N}(10^{-2}, 10^{-8})$; $W_{\mathrm{CA1}} \sim \mathcal{N}(10^{-2}, 10^{-8})$; $W_{\mathrm{pred}} \leftarrow 0$
    $\mathcal{A}_{\mathrm{CA3}} \leftarrow 0$; $\mathrm{ET} \leftarrow 0$; $\mathrm{IS}_{\mathrm{CA3}} \leftarrow 0$; $\mathrm{IS}_{\mathrm{CA1}} \leftarrow 0$

**for** lap $= 1$ **to** TotalLaps **do**
    **for** $s \in \mathcal{S}_{\mathrm{lap}}$ **do**
        **for** $t = 1$ **to** $T_s$ **do**
            `# Generate new spike trains and calculate the activities at every` $T_{\mathrm{SpikeUpdate}}$.
            **if** $\mathrm{mod}(t, T_{\mathrm{SpikeUpdate}}) = 0$ **then**
                $\mathrm{SpikeTrain}_{\mathrm{CA3}} \leftarrow \mathrm{GenerateSpikeTrain}(s, \mathcal{A}_{\mathrm{CA3}}, W_{\mathrm{CA3}})$
                $\mathcal{A}_{CA3} \leftarrow \mathrm{Activity}(\mathrm{SpikeTrain}_{\mathrm{CA3}})$

                $\mathrm{SpikeTrain}_{\mathrm{CA1}} \leftarrow \mathrm{GenerateSpikeTrain}(\cdot, \mathcal{A}_{\mathrm{CA3}}, W_{\mathrm{CA1}})$
                $\mathcal{A}_{CA1} \leftarrow \mathrm{Activity}(\mathrm{SpikeTrain}_{\mathrm{CA1}})$
            **end**

            `# Compute novelty and train the feature prediction network.`
            $(N(t), W_{\mathrm{pred}}) \leftarrow \mathrm{DeltaRule}(P_s, \mathcal{A}_{\mathrm{CA1}}, W_{\mathrm{pred}})$
            $\mathrm{PS}(t) \leftarrow \mathrm{PerceivedSalience}(I_s, N(t), \mathrm{MI})$

            `# Calculate the plateau probability of hippocampal neurons.`
            $p_{\mathrm{plataeu,CA3}} \leftarrow \mathrm{PlateauProbability}(\mathcal{A}_{\mathrm{CA3}}, \cdot)$
            $p_{\mathrm{plataeu,CA1}} \leftarrow \mathrm{PlateauProbability}(\mathcal{A}_{\mathrm{CA1}}, \mathrm{PS}(t))$

            `# Update ETs and ISs.`
            $\mathrm{ET} \mathrel{+}= \mathrm{UpdateET}(\mathrm{SpikeTrain}_{\mathrm{CA3}})$
            $\mathrm{IS}_{\mathrm{CA3}} \mathrel{+}= \mathrm{UpdateIS}(p_{\mathrm{plataeu,CA3}})$
            $\mathrm{IS}_{\mathrm{CA1}} \mathrel{+}= \mathrm{UpdateIS}(p_{\mathrm{plataeu,CA1}})$

            `# Train the hippocampal network.`
            $W_{\mathrm{CA3}} \leftarrow \mathrm{BTSP}(\mathrm{ET}, \mathrm{IS}_{\mathrm{CA3}}, W_{\mathrm{CA3}})$
            $W_{\mathrm{CA1}} \leftarrow \mathrm{BTSP}(\mathrm{ET}, \mathrm{IS}_{\mathrm{CA1}}, W_{\mathrm{CA1}})$

            `# Decay eligibility traces and instructive signals.`
            $\mathrm{ET} \mathrel{-}= \mathrm{ET}/\tau_{\mathrm{ET}}$
            $\mathrm{IS}_{\mathrm{CA3}} \mathrel{-}= \mathrm{IS}_{\mathrm{CA3}}/\tau_{\mathrm{IS}_{\mathrm{CA3}}}$
            $\mathrm{IS}_{\mathrm{CA1}} \mathrel{-}= \mathrm{IS}_{\mathrm{CA1}}/\tau_{\mathrm{IS}_{\mathrm{CA1}}}$
        **end**
    **end**
**end**

### 3 Offline phase simulation details

Offline-phase simulations were performed using python library Brian 2 [44], using the synaptic weight matrices $W_{\text{CA3}}$ and $W_{\text{CA1}}$ obtained from the online learning phase. Starting from these learned weights, we adopted the offline simulation framework of Ecker et al.[11], using their publicly available code as the basis for our implementation.

#### 3.1 Single cell model

We used a neuron model with the adaptive exponential integrate-and-fire (AdExIF) equations [35, 15], as in Ecker et. al [11]. Here, the membrane potential $V(t)$ and adaptation current $w(t)$ interact, following the equations:

$$C_m \frac{dV(t)}{dt} = -g_L(V(t) - V_{\text{rest}}) + g_L \Delta_T \exp\left(\frac{V(t) - \vartheta}{\Delta_T}\right) - w(t) + I_{\text{syn}}(t), \tag{1}$$

$$\tau_w \frac{dw(t)}{dt} = a(V(t) - V_{\text{rest}}) - w(t). \tag{2}$$

When $V(t)$ crosses the threshold $\vartheta$, it is reset to $V_{\text{reset}}$ and held for a refractory period $t_{\text{ref}}$. The model incorporates adaptation process, which is important in replay generation [11]. The adaptation current $w(t)$ is incremented by $b$ at each spike, where $a$ and $b$ control subthreshold and spike-triggered adaptation, respectively.

The actual parameter values used for the simulations are summarized in Table 1, adapted from the previous work [11]. Physical units are as follows: $C_m$ (pF), $g_L$ and $a$ (nS), $V_{\text{rest}}$, $\Delta_T$, $\vartheta$, and $V_{\text{reset}}$ (mV), $t_{\text{ref}}$ and $\tau_w$ (ms), and $b$ (pA).

Table 1: Neuron model parameters.

| Parameter | Symbol | Value |
|---|---|---|
| Membrane capacitance | $C_m$ | 180.13 |
| Leak conductance | $g_L$ | 5 |
| Leak reversal (resting) potential | $V_{\text{rest}}$ | -75.19 |
| Slope of exponential term | $\Delta_T$ | 4.23 |
| Firing threshold | $\vartheta$ | -24.42 |
| Adaptation time constant | $\tau_w$ | 84.93 |
| Subthreshold adaptation strength | $a$ | -0.01 |
| Spike-triggered adaptation increment | $b$ | 500 |
| Reset potential | $V_{\text{reset}}$ | -29.74 |
| Absolute refractory period | $t_{\text{ref}}$ | 5.96 |

#### 3.2 Synapse model

Synapses were modeled as conductances with biexponential kinetics:

$$g(t) = \hat{g}A\left[\exp\left(-\frac{t}{\tau_d}\right) - \exp\left(-\frac{t}{\tau_r}\right)\right], \tag{3}$$

where $\hat{g}$ is the peak conductance, and $\tau_r$ and $\tau_d$ are the rise and decay time constants, respectively.

The normalization constant $A = \exp(-t_p/\tau_d) - \exp(-t_p/\tau_r)$ ensures that the conductance peaks at time $t_p = \tau_r \tau_d/(\tau_d - \tau_r)\ln(\tau_d/\tau_r)$. Synaptic currents were computed as

$$I_{\text{syn}}(t) = g_{\text{AMPA}}(t)\,(V(t) - E_{\text{exc}}) + g_{\text{GABA}}(t)\,(V(t) - E_{\text{inh}}), \tag{4}$$

where $E_{\text{exc}} = 0$ mV and $E_{\text{inh}} = -70$ mV are the reversal potentials of excitatory and inhibitory synapses, respectively.

The parameters used for the synapses are summarized in Table 2. Ext stands for excitatory cells, Inh stands for inhibitory cells, and EC stands for entorhinal cortex populations. Physical dimensions are as follows: $\hat{g}$ (nS), $\tau_r$, $\tau_d$, and $t_d$ (ms), and connection probability $p_{\text{conn}}$ is dimensionless.

Table 2: Synaptic parameters (adapted from [11]).

| **Connection** | $\hat{g}$ (nS) | $\tau_r$ (ms) | $\tau_d$ (ms) | $t_d$ (ms) | $p_{\text{conn}}$ |
|---|---|---|---|---|---|
| Ext → Ext | 0.1-6.3 | 0-15 | 1.3 | 9.5 | 0.1 |
| Ext → Inh | 0.85 | 1.0 | 4.1 | 0.9 | 0.1 |
| Inh → Ext | 0.65 | 0.3 | 3.3 | 1.1 | 0.25 |
| Inh → Inh | 5.0 | 0.25 | 1.2 | 0.6 | 0.25 |
| EC → Ext | 19.15 | 21.5 | 5.4 | – | – |

#### 3.3 Connection weight optimization

Synaptic weights of the network were optimized using the publicly available code from the previous study [11] after being modified to incorporate $W_{\text{CA1}}$ network. We sought to determine an optimal set of single-valued parameters determining the strength of excitatory and inhibitory connections within and between hippocampal subfields. Specifically, the parameters optimized are listed below:

- Entorhinal cortex (EC) input rate
- EC → CA3 excitatory connection weight
- CA3 excitatory → CA3 inhibitory connection weight
- CA3 inhibitory → CA3 excitatory connection weight
- CA3 inhibitory → CA3 inhibitory connection weight
- Weight scaling factor for $W_{\text{CA3}}$
- CA1 excitatory → CA1 inhibitory connection weight
- CA1 inhibitory → CA1 excitatory connection weight
- CA1 inhibitory → CA1 inhibitory connection weight

Optimization followed the evolutionary multi-objective framework [11], with customized upper and lower bounds for each parameter listed in Table 3. The final optimized weights used in the three task environments are summarized in Table 4.

Table 3: Upper bound and lower bound values used for the optimization process.

| Parameter | Lower bound | Upper bound |
|---|---|---|
| EC input rate | 5.0 | 15.0 |
| EC → CA3 Ext | 0.5 | 40.0 |
| CA3 Exc → Inh | 0.2 | 5.0 |
| CA3 Inh → Exc | 0.5 | 20.0 |
| CA3 Inh → Inh | 1.0 | 15.0 |
| CA3 weight scale ($W_{\text{CA3}}$) | 1.0 | 2.5 |
| CA1 Exc → Inh | 0.2 | 5.0 |
| CA1 Inh → Exc | 1.0 | 10.0 |
| CA1 Inh → Inh | 1.0 | 15.0 |

Table 4: Optimized connection weights. Each value represents the final parameter obtained after evolutionary optimization.

| Parameter | Linear treadmill | T-maze | Closed linear track |
|---|---|---|---|
| EC input rate | 10.084 | 7.322 | 13.038 |
| EC → CA3 Ext | 23.957 | 34.481 | 15.843 |
| CA3 Exc → Inh | 0.349 | 1.765 | 1.715 |
| CA3 Inh → Exc | 1.941 | 1.381 | 1.646 |
| CA3 Inh → Inh | 2.971 | 11.110 | 14.732 |
| CA3 weight scale ($W_{\text{CA3}}$) | 2.210 | 1.886 | 2.306 |
| CA1 Exc → Inh | 1.000 | 1.765 | 0.337 |
| CA1 Inh → Exc | 1.198 | 1.877 | 1.052 |
| CA1 Inh → Inh | 14.725 | 13.819 | 12.671 |

##### 3.4 Position decoding and replay event detection

Using the simulated spontaneous activity of CA3 neurons during the offline phase, we detected individual replay events following the spatial position decoding procedure described by Davidson et al. [9].

We first identified candidate replay periods as time blocks in which the mean firing rate of the CA3 population exceeded 2 Hz for at least 200 ms. Only these high-activity periods were considered for subsequent decoding analysis.

For each candidate event, we defined a set of possible decoded positions along the trajectory of interest. These positions were defined by subdividing each behavioral state into four equally spaced spatial points. For example, when analyzing shortcut replay events (Fig. 5), states 4–10 were included, resulting in a total of 28 candidate spatial positions.

CA3 firing rates were computed in sliding 10 ms time windows across the candidate replay event. For each time bin, we estimated the posterior probability of the animal being at each candidate position by comparing the observed CA3 firing rates with the ideal spatial tuning curves of each CA3 neuron (see Methods). As a result, a posterior probability matrix $X_{\text{posterior}}$ was obtained, indexed by spatial position and time.

To quantify whether the decoded activity formed a coherent sequential trajectory, we fitted a straight line to the posterior matrix, assuming a constant-velocity traversal through space. The ”goodness” of fit of this line was quantified using the metric $R$, adapted from Davidson et al. [9]. Specifically, $R$ measures the fraction of posterior probability mass being included within a narrow band surrounding the fitted line:

$$R = \frac{\sum_{x,t} X_{\text{posterior}}(x,t)\,\mathbb{I}\big(|x-\hat{x}(t)| < d\big)}{\sum_{x,t} X_{\text{posterior}}(x,t)},$$

where $\hat{x}(t)$ denotes the position predicted by the fitted line at time $t$, $\mathbb{I}$ is an indicator function, and $d$ is a spatial tolerance (band size). To ensure that the fitted trajectory spanned a meaningful portion of the spatial domain, we required that the fitted line pass through at least one-third of the spatial bins; otherwise, $R$ was set to zero.

To assess statistical significance, we generated a null distribution by repeating the same decoding procedure 100 times using shuffled CA3 spatial tuning curves. A candidate event was classified as a valid replay if its true $R$ value exceeded the 95th percentile of the shuffled $R$ distribution. The slope and intercept of the fitted line were then used to classify replay direction and trajectory type.

### 4 Supplementary figures

#### 4.1 Details of synaptic learning rule during the online phase

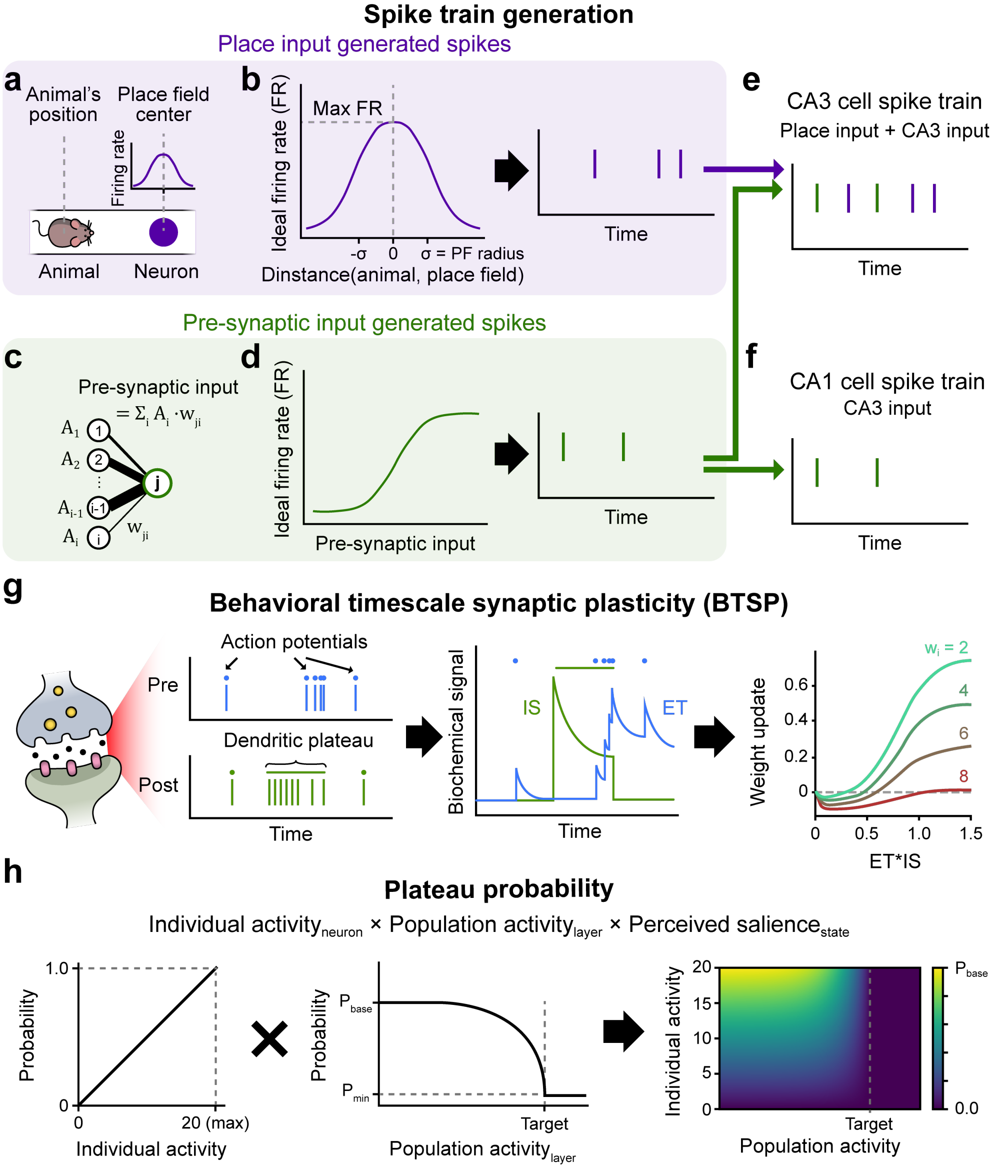

**Supplementary Figure 1.** Spiking neural network model activity. **a–f**, Spike-generation process. **a**, Place-input–driven firing rates were determined by the distance between the animal and each place cell's field center. **b**, This distance was converted to a firing rate using a Gaussian tuning curve. **c**, Pre-synaptic input–driven firing rates were computed from the weighted sum of pre-synaptic activity. **d**, The weighted sum was passed through a sigmoid nonlinearity to obtain the final rate. **e**, CA3 cell firing rates were computed as the sum of place-input–generated and pre-synaptic–input–generated components. **f**, CA1 cell firing rates were determined solely by pre-synaptic–input–generated activity. **g**, Behavioral time-scale synaptic plasticity (BTSP). Each pre-synaptic action potential produced an eligibility trace (ET), and each post-synaptic dendritic plateau potential (PP) generated an instructive signal (IS). Synaptic weight updates depended on the product of ET and IS. **h**, Modulation of PP probability. In our model, the probability of PP occurrence was computed as the product of a neuron's recent activity, population-level normalization, and the perceived salience of the current state.

#### 4.2 Online phase simulation using symmetric STDP

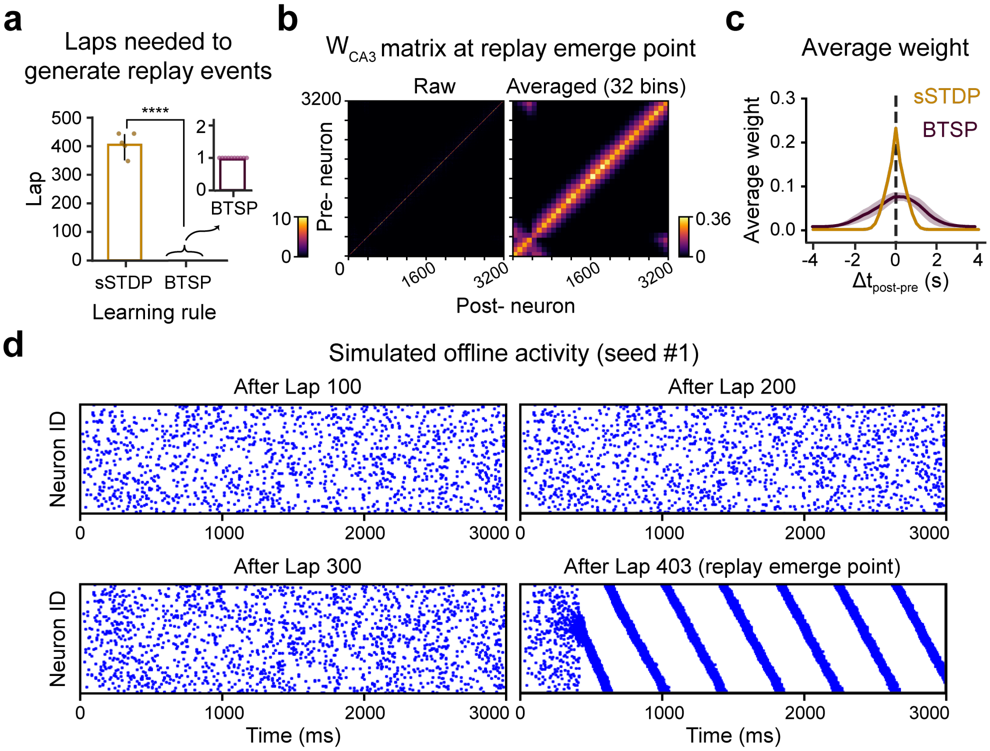

**Supplementary Figure 2.** Laps the animal needs to travel for its CA3 network to acquire spontaneously-generated reply activity, in the linear-treadmill simulations. **a**, Replay emerge point, defined by the number of laps needed to generate spontaneous replays. The networks trained with symmetric STDP (sSTDP) demonstrated much later emergence of replay, compared to one-shot acquisition of replay within the networks trained using BTSP. **b**, $W_{CA3}$ of a network trained using sSTDP at the replay emerge point. Left: $W_{CA3}$, Right: Averaged $W_{CA3}$ computed by dividing the matrix into 32 bins and computed, plotted averaged value of each bin. **c**, Average connection weight profile of networks trained by sSTDP and BTSP. sSTDP trained weights showed stronger connection weight to other CA3 place cells with nearby place fields. BTSP trained weights are projected more broadly, projecting weights to place cells with far place fields and less strong connections to cells with nearby place fields. **d**, Simulated offline phase of sSTDP trained network. Before the replay emerge point, the activity does not show significant clustered firing or replay like activity. At lap 403 which is its replay emerge point, replay activity emerge.

#### 4.3 Equilibrium properties of BTSP

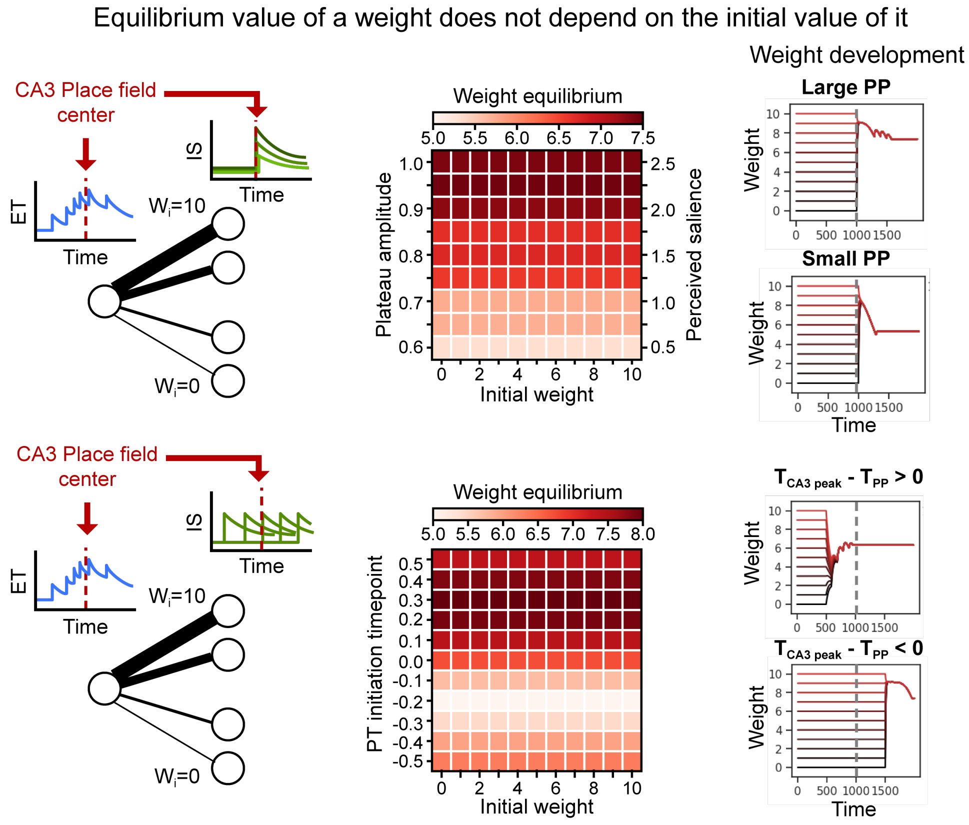

**Supplementary Figure 3.** Simulations of synaptic weight updates as a function of plateau amplitude (top) and plateau onset timing (bottom), across different initial synaptic weight strengths. The results show that the final synaptic weights are modulated by plateau amplitude and onset timing, but not by the initial synaptic weight value.

#### 4.4 Implementing layerwise feedback inhibition during place cell activity estimation

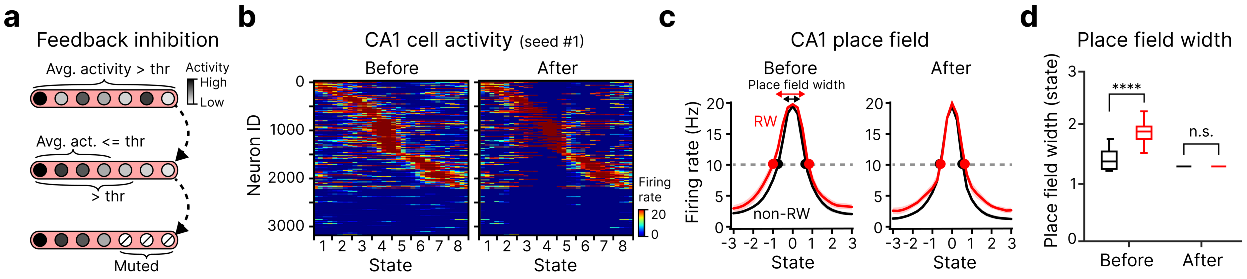

**Supplementary Figure 4.** Implementing layerwise feedback inhibition during place cell activity estimation. **a**, Schematic of layerwise feedback inhibition. Top: identify positions at which the average CA1 population activity exceeds a predefined threshold. Middle: select the maximal subset of neurons, starting from the most active, such that the average activity does not exceed the threshold. Bottom: selectively suppress the remaining less active neurons, reducing the overall population activity to the target level. **b**, Example CA1 cell activity before (left) and after (right) applying feedback inhibition. Activity after lap 10 from Fig. 3 is shown, where a reward is introduced at state 4. **c**, Average place-field tuning curves before (left) and after (right) feedback inhibition. Place cells encoding reward-associated locations (red) and non-reward locations (black) are shown separately. The dashed grey line indicates the firing-rate threshold used to define active place fields (15 Hz). Data are presented as mean±s.d. **d**, Place-field width before (left) and after (right) feedback inhibition. Place-field width was defined as the spatial extent over which firing rates exceeded the activity threshold. Incorporating feedback inhibition eliminates reward-induced place-field broadening. Independent t-test, $N_{\text{seed}} = 10$, ****$P = 3.47 \times 10^{-5}$; n.s., $P = 1.0$.

#### 4.5 Replay frequency modulation based on perceived salience

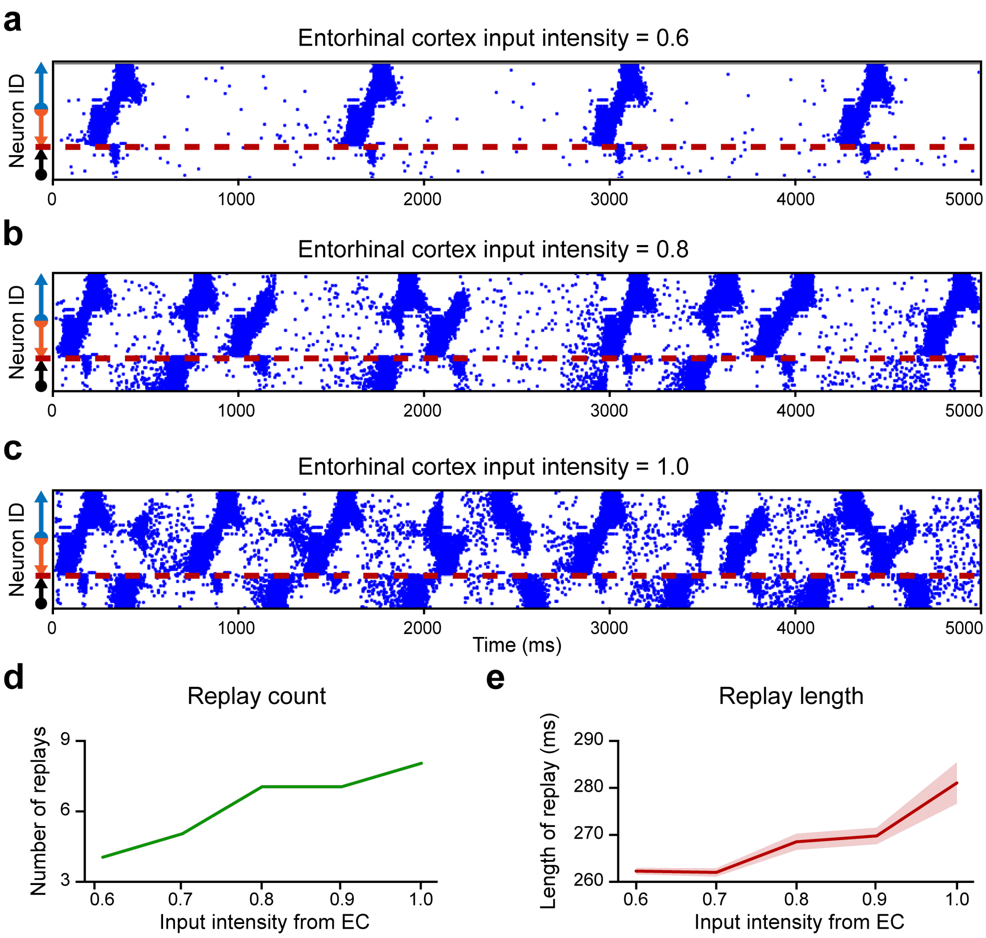

**Supplementary Figure 5.** Replay frequency and event length are modulated by the strength of entorhinal cortex (EC) input. **a–c**, Simulated offline activity in networks that learned the T-maze task under different EC input strengths (related to Fig. 4–5). **a**, EC input at 60% of the standard strength. **b**, EC input at 80% of the standard strength. **c**, EC input at 100% (full) strength. **d**, Number of replay events as a function of EC input strength. **e**, Replay event length as a function of EC input strength.

[39] Rajat Saxena, Justin L Shobe, and Bruce L McNaughton. “Learning in deep neural networks and brains with similarity-weighted interleaved learning”. en. In: *Proc. Natl. Acad. Sci. U. S. A.* 119.27 (July 2022), e2115229119.

[52] Zhenglong Zhou, Michael J Kahana, and Anna C Schapiro. “A unifying account of replay as context-driven memory reactivation”. Mar. 2023.
